# Combining supervised and unsupervised analyses to quantify behavioral phenotypes and validate therapeutic efficacy in a triple transgenic mouse model of Alzheimer’s disease

**DOI:** 10.1101/2024.06.07.597924

**Authors:** Thais Del Rosario Hernandez, Narendra R. Joshi, Sayali V. Gore, Jill A. Kreiling, Robbert Creton

**Affiliations:** Department of Molecular Biology, Cell Biology and Biochemistry, Brown University, Providence, Rhode Island, United States

## Abstract

Behavioral testing is an essential tool for evaluating cognitive function and dysfunction in preclinical research models. This is of special importance in the study of neurological disorders such as Alzheimer’s disease. However, the reproducibility of classic behavioral assays is frequently compromised by interstudy variation, leading to ambiguous conclusions about the behavioral markers characterizing the disease. Here, we identify age- and genotype-driven differences between 3xTg-AD and non-transgenic control mice using a low-cost, highly customizable behavioral assay that requires little human intervention. Through behavioral phenotyping combining both supervised and unsupervised behavioral classification methods, we are able to validate the preventative effects of the immunosuppressant cyclosporine A in a rodent model of Alzheimer’s disease, as well as the partially ameliorating effects of candidate drugs nebivolol and cabozantinib.

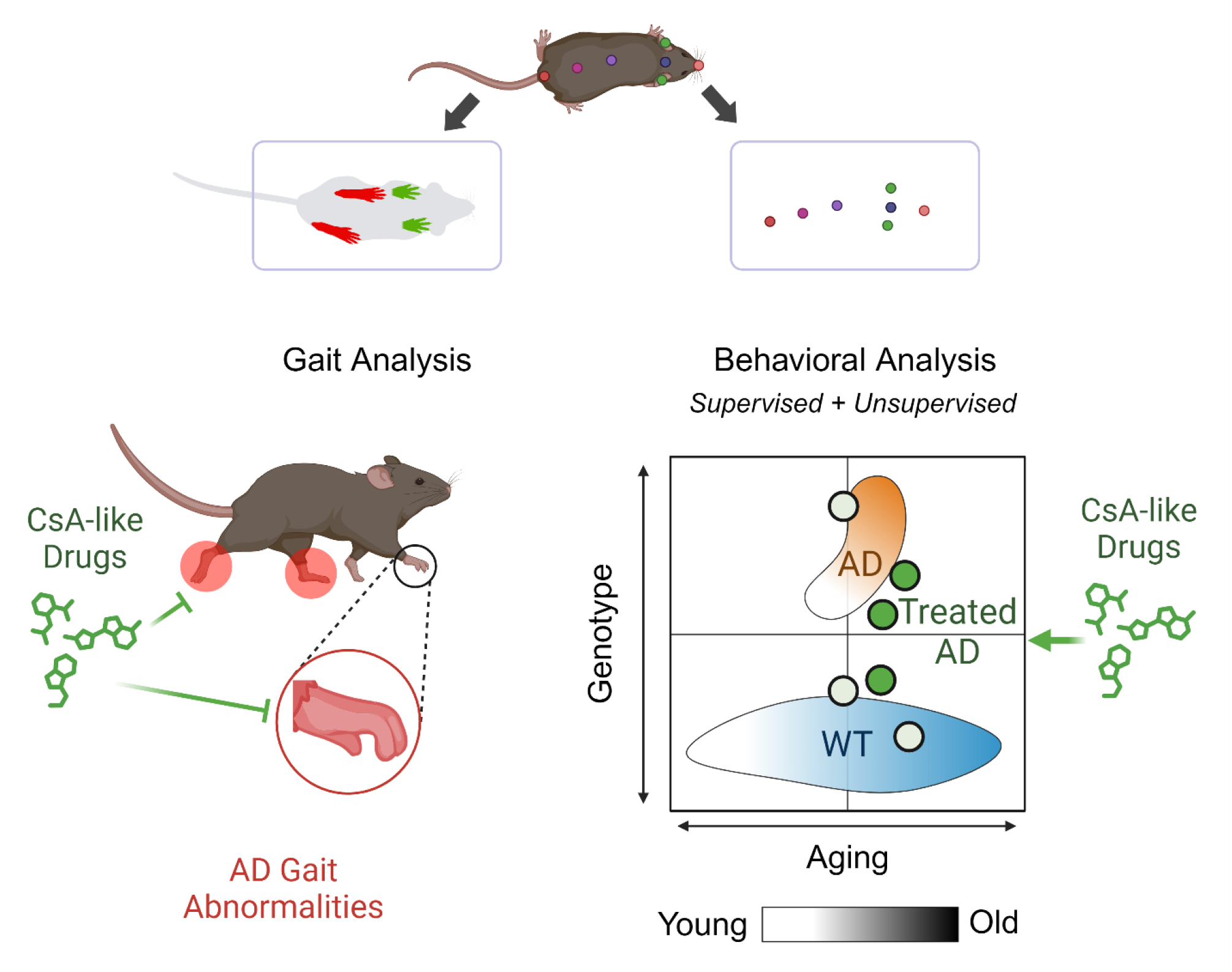

## Introduction

Alzheimer’s disease (AD) is a neurodegenerative disorder characterized by progressive cognitive decline caused by the accumulation of amyloid-beta plaques, neurofibrillary tangles, and synaptic dysfunction^1^. Despite extensive research efforts, the detection and potential prevention of early AD pathology continues to be a challenging task, further complicated by the slow and complex onset of the disease^2^. A promising candidate therapeutic is cyclosporine A (CsA), an immunosuppressant drug shown to reduce the incidence of AD in transplant patients^3,4^. While CsA is not an ideal candidate drug due to the side effects caused by its immunosuppressive pharmacological mechanism, we hypothesized that CsA-like drugs might have the same neuroprotective effects with reduced side effects. In a previous study using high-throughput screening in zebrafish larvae, we were able to identify several CsA-like compounds with potential neuroprotective effects, including the beta-adrenergic antagonist nebivolol and the non-specific tyrosine kinase inhibitor cabozantinib^5,6^. To expand on this work, our current study looks at the efficacy of these drugs in a mammalian model of AD.

Many available mouse models of AD have been well characterized and are used to study different components of AD pathology^7^. In this study, we focused on 3xTg-AD mice, a triple transgenic mouse model harboring APP^Swe^ and tau^P301L^ transgenes on a mutant PS1^M146V^ knock-in background^8^. This well-established model has been shown to exhibit early molecular and behavioral markers of AD pathology, making it a suitable model for the evaluation of preventive treatments^9,10^. Moreover, several studies performed on 3xTg-AD mice focus on disease-relevant parameters such as spatial learning and memory, spontaneous alternation behavior, and associative learning^9,11^. However, this mouse model is not exempt from the interstudy variation of behavioral phenotypes that is commonly found in preclinical research involving animal models of multifaceted neurodegenerative diseases. Similarly to other AD models, common behavioral assays such as the elevated plus maze, open-field test, and Y-maze often exhibit study-dependent results in 3xTg-AD mice^12-15^. This abundance of conflicting results highlights the prevalent challenges of behavioral characterization in AD mouse models.

Due to the complex and gradual progression of AD, there is a growing need for robust and comprehensive behavioral assays that can adequately capture behavioral phenotypes in high resolution and across timepoints - especially those that are relevant to early disease states. We have previously developed a highly automated and noninvasive multi-animal behavioral analysis pipeline capable of measuring subtle changes in mouse behavior within a familiar environment^16^. In the current study, we propose a multifaceted, longitudinal behavioral assessment to 1) validate the effect of chronic CsA treatment in a triple transgenic AD mouse model, 2) determine the efficacy of previously identified CsA-like drugs in ameliorating AD pathology, and 3) establish a novel behavioral assay capable of detecting both early and late disease stage phenotypic markers. We posit that the quantification of behavioral parameters with supervised and unsupervised learning methods will greatly enhance the sensitivity and depth of AD-relevant behaviors.

## Results

### Comparing gait alterations between WT and 3xTg-AD mice across stages of aging

We carried out detailed gait analysis in 7-month-old (“young”) and 14-month-old (“old”) 3xTg-AD and non-transgenic control mice (B6129SF2/J, henceforth referred to as WT mice) using the Noldus CatWalk XT system. Mice were placed on the system’s glass walkway and allowed to walk from one end to another. Each run was recorded using a high-resolution camera, included with the system. Animals were recorded for at least 3 compliant runs, and individual paws were classified automatically using the CatWalk software (RF: right forepaw; LF: left forepaw; RH; right hindpaw; LH: left hindpaw) (Figure 1A). We measured 10 paw parameters, 2 support parameters, and 4 step parameters (Table 1). We found that several parameters showed significant differences between WT and 3xTg-AD mice.

**Table 1.** Gait parameters measured with CatWalk XT.

|  | **CatWalk XT Measure** | **Description** |
| --- | --- | --- |
| **Paw Parameters** | Manual print length | Distance between the third toe and the heel of a paw. |
|  | Toe spread | Distance between the first and fifth toe of a hind paw. |
|  | Intermediate toe spread | Distance between the second and fourth toe of a paw. |
|  | Print area | Average surface area of the paw print. |
|  | Maximum contact area | Surface area of a print when it makes maximum contact with the glass walkway. |
|  | Mean intensity | Average intensity of a print. |
|  | Minimum intensity | Minimum intensity of a print. |
|  | Stand | Duration of contact of a print with the glass walkway. |
|  | Paw angle body axis | Angle of the paw relative to the orientation of the body axis of the animal. |
|  | Step cycle | Time between two consecutive initial contacts of the same paw. |
| **Support Parameters** | Support | Relative duration of simultaneous contact with the glass walkway of different paw combinations - namely, girdle paws (RF-LF or RH-LH), lateral paws (RF-RH or LF-LH), diagonal (RF-LH or LF-RH), all four paws, three paws, a single paw, and zero paws. |
|  | Base of support | Average width between the front or hind paws. |
| **Step Parameters** | Duration | Duration of the run based on the identified steps. |
|  | Cadence | Number of steps per second. |
|  | Number of steps | Total number of identified steps in a run. |
|  | Step sequence (% of NSSP) | Percentage of time spent in one of the six normal step sequence patterns. |

**Figure 1.**
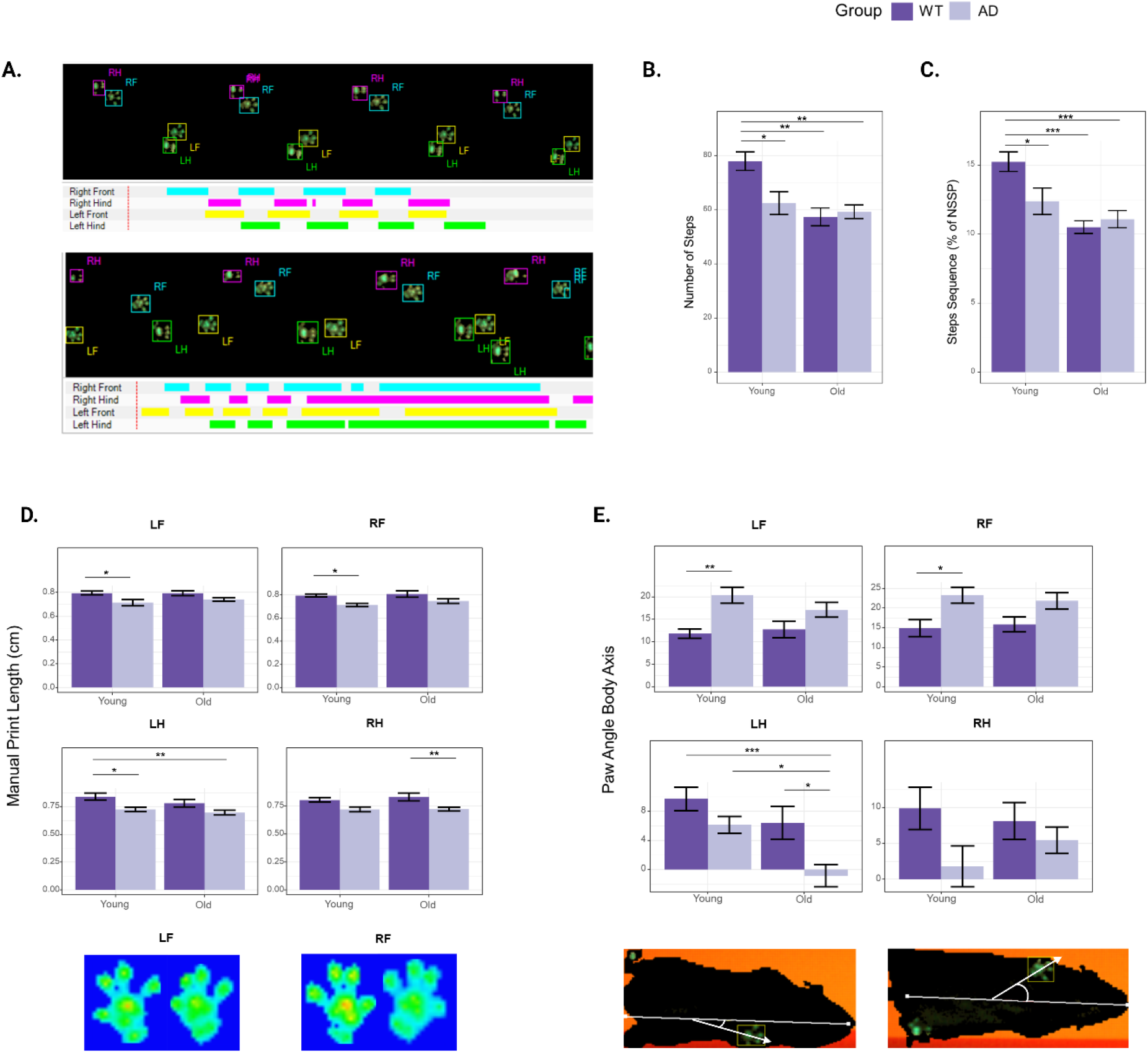
Gait analysis of 3xTg-AD and WT mice. A) Example of WT (top) and 3xTg-AD (bottom) paw prints automatically recorded and categorized with the CatWalk XT system. B) Number of classified steps. C) Percentage of time spent in a normal step sequence pattern (NSSP). D) Manual print length, E) intermediate toe spread, F) mean intensity, and G) paw angle body axis for all four paws. The results are expressed as mean ± SEM. Statistics: two-way ANOVA followed by post hoc Tukey HSD test, * p < 0.05, ** p < 0.01, *** p < 0.001. N = 8 WT mice, N = 9 3xTg-AD mice.

The number of steps was significantly higher in young WT mice compared to young 3xTg-AD mice (*p* = 0.016). The number of steps significantly decreased with age in WT mice (*p =* 0.001) but not in 3xTg-AD mice (*p* = 0.891) (Figure 1B). Similarly, the percentage of normal step sequence patterns (NSSP) was significantly different between young WT and young 3xTg-AD mice (*p* = 0.044), as well as between young and old WT mice (*p* = 0.0004), but not between young and old 3xTg-AD mice (p = 0.552) (Figure 1C). These results suggest that the number of steps and the percentage of NSSP are both parameters that decrease with normal aging but are already in deficit in young 3xTg-AD mice. Next, we looked at paw-specific parameters and found significant differences in print length (LF: *p* = 0.032, RF: *p* = 0.048) and paw angle body axis (LF: *p* = 0.007, RF: *p* = 0.049) of the front paws of young WT and young 3xTg-AD mice. As the mice aged, these differences became non-significant (Figure 1D, 1E). These parameters identify early gait variations in 3xTg-AD mice. In contrast, quantification of both mean intensity and minimum intensity of the front paws reveal significant differences only between old 3xTg-AD mice and WT mice. However, this might be attributed to the difference in weight between the genotypes (Figure 3C).

Interestingly, we found that 3xTg-AD mice place more weight on their hind limbs as they age, as measured by the base of support parameter (*p* = 0.017). This increase was also captured in the RH measurements for the stand (*p* = 0.016) and step cycle parameters (*p* = 0.024), which measure the amount of time a paw is in contact with the glass walkway and the time between two consecutive paw contacts, respectively. Additionally, we observed a significant increase in the mean intensity (LH: *p* = 0.002, RH: *p* = 0.00001) and minimum intensity (LH: *p* = 0.007, RH: *p* = 0.006) of both hind paws in 3xTg-AD mice, but no significant changes in WT mice as they age. The statistical significance analysis for all gait parameters measured in this study is provided in Supplementary Table 1.

### Detecting differences across age and genotype using automated 8-cage analysis

In order to characterize the behavioral phenotype of the triple transgenic Alzheimer’s disease mouse model (3xTg-AD), we performed an automated analysis of behavior in our 8-cage imaging system (Figure 2A). Mice are presented with a 22-hour PowerPoint simulating daytime and nighttime, as well as two stimuli: a spinning yellow moth, and red moving lines in the home quadrant (Figure 2B). We compared them to age-matched, non-transgenic controls (WT) across three timepoints: young (5-6 months), old (12-13 months), and aged (18-19 months). Data from the acquired videos was analyzed using Fiji, and 15 behavioral parameters measuring movement, acclimation, habituation, and time spent in each quadrant of the cage were quantified (Table 2). 4 of these 15 behaviors were significantly different between WT and 3xTg-AD mice in at least one of the age timepoints - namely, movement during the 1st hour (*p* = 0.001), movement during the first hour of nighttime (young: *p* = 0.021, old: *p* = 0.034), acclimation to the cage in % stretch-attend posture (SAP) (*p* = 0.029), and habituation to the moth stimulus in % moved (*p* = 0.018) (Figure 2E).

**Table 2.** Behavioral parameters measured using Fiji.

| Behavioral measure | Description |
| --- | --- |
| M1 | Movement during the first hour |
| MD | Movement during daytime |
| M7 | Movement in the 7<sup>th</sup> hour (i.e. the first hour of nighttime) |
| MN | Movement during nighttime |
| N-D | The difference between nighttime and daytime movement (MN-MD) |
| AM | Acclimation to the cage, measured in % movement |
| AS | Acclimation to the cage, measured in % stretch-attend posture (SAP) |
| HMM | Habituation to moth stimulus, measured in % movement |
| HL1M | Habituation to 1<sup>st</sup> set of moving lines, measured in % movement |
| HL2M | Habituation to 2<sup>nd</sup> set of moving lines, measured in % movement |
| Home | % time the mouse is in the home quadrant |
| Wall | % time the mouse is in the wall quadrant |
| Food | % time the mouse is in the food quadrant |
| Window | % time the mouse is in the window quadrant |
| Out | Sum of Wall and Window. |

**Figure 2.**
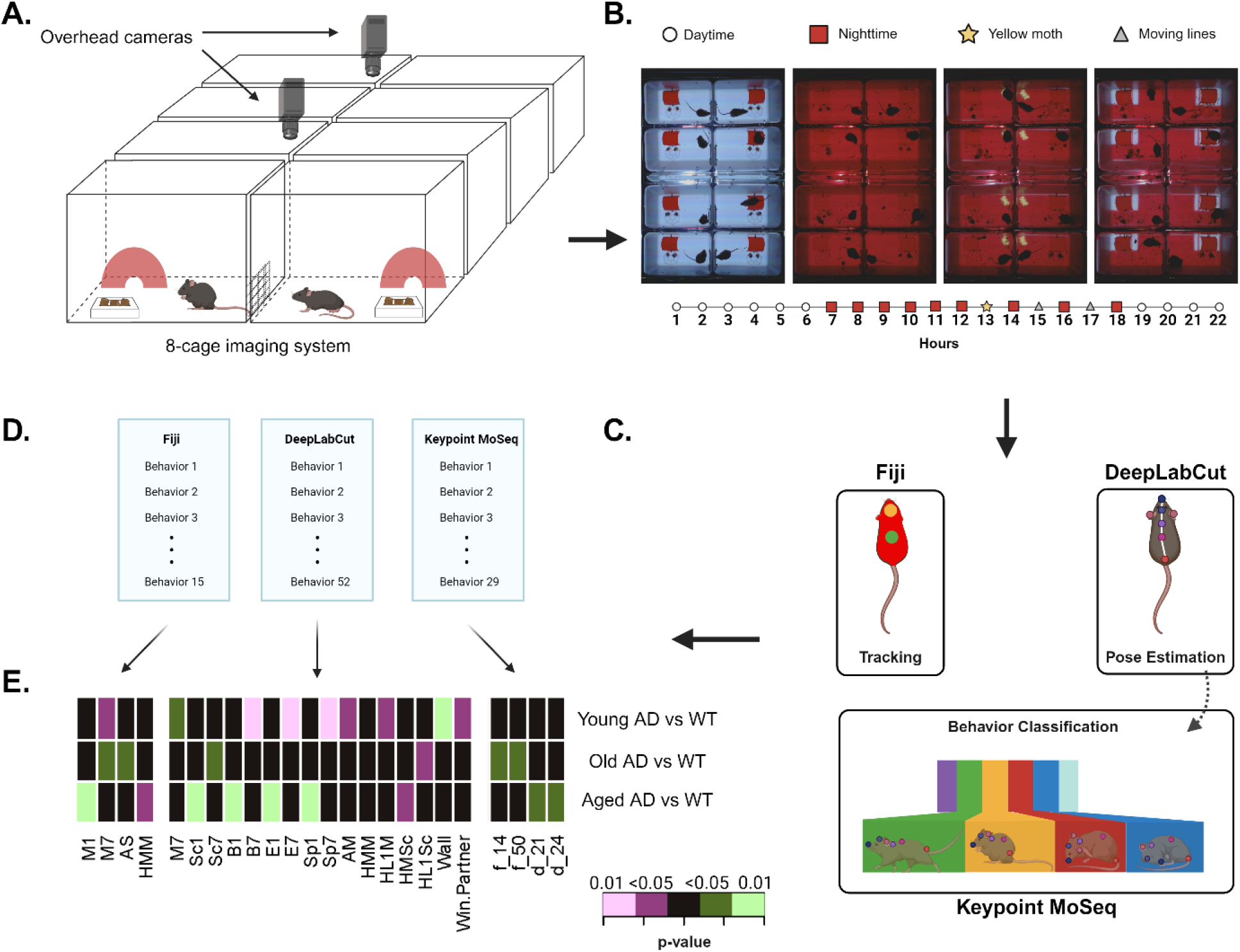
Overview of 8-cage imaging system and analysis pipeline. A) Mice are individually placed in a cage with access to food, water, and shelter in the form of a red hut. B) They are recorded for 22 hours, during which they are shown stimuli three times (hours 13, 15, and 17). C) The videos are analyzed using Fiji, DeepLabCut, and Keypoint MoSeq and D) A total of 125 behaviors are extracted. E) Significantly different behaviors in 3xTg-AD mice compared to age-matched WT mice. Green indicates an increase in 3xTg-AD mice compared to WT mice, while purple indicates a decrease. Statistics: Two-sample Wilcoxon test, * p < 0.05, ** p < 0.01. N = 8 WT mice, N = 16-18 3xTg-AD mice.

We also analyzed the videos using our previously trained DeepLabCut (DLC) model to identify an additional 52 behavioral parameters (Table 3). Of these, 15 showed significant differences between WT and 3xTg-AD mice during one of the age timepoints: movement, scoot, burst, escape, and speed during the first hour of nighttime (*p* = 0.048, *p* = 0.025, *p* = 0.002, *p* = 0.003, and *p* = 0.008, respectively), scoot, burst, escape, and speed during the first hour (*p* = 0.001, *p* = 0.001, *p* = 0.001, and *p* = 0.001, respectively), acclimation measured with % movement (*p* = 0.032), habituation to moth stimulus in % scoot (*p* = 0.040), habituation to the first set of moving lines in % movement and % scoot (*p* = 0.021 and *p* = 0.043, respectively), distance from any cage wall (*p* = 0.001), and time spent near the window with partner (*p* = 0.028).

**Table 3.** Behavioral parameters measured using DeepLabCut.

| Behavioral measure | Description |
| --- | --- |
| M1 | Movement during the first hour |
| MD | Movement during daytime |
| M7 | Movement in the 7<sup>th</sup> hour (i.e. the first hour of nighttime) |
| MN | Movement during nighttime |
| MN-D | The difference between nighttime and daytime movement (MN-MD) |
| Sc1 | Scoot during the first hour |
| ScD | Scoot during daytime |
| Sc7 | Scoot in the 7<sup>th</sup> hour (i.e. the first hour of nighttime) |
| ScN | Scoot during nighttime |
| ScN-D | The difference between nighttime and daytime scoot (ScN-ScD) |
| B1 | Burst during the first hour |
| BD | Burst during daytime |
| B7 | Burst in the 7<sup>th</sup> hour (i.e. the first hour of nighttime) |
| BN | Burst during nighttime |
| BN-D | The difference between nighttime and daytime burst (BN-BD) |
| E1 | Escape during the first hour |
| ED | Escape during daytime |
| E7 | Escape in the 7<sup>th</sup> hour (i.e. the first hour of nighttime) |
| EN | Escape during nighttime |
| EN-D | The difference between nighttime and daytime escape (EN-ED) |
| Sp1 | Speed during the first hour |
| SpD | Speed during daytime |
| Sp7 | Speed in the 7<sup>th</sup> hour (i.e. the first hour of nighttime) |
| SpN | Speed during nighttime |
| SpN-D | The difference between nighttime and daytime speed (SpN-SpD) |
| AM | Acclimation to the cage, measured in % movement |
| ASc | Acclimation to the cage, measured in % scoot |
| AB | Acclimation to the cage, measured in % burst |
| AE | Acclimation to the cage, measured in % escape |
| ASp | Acclimation to the cage, measured in speed (pixels) |
| HMM | Habituation to moth stimulus, measured in % movement |
| HL1M | Habituation to 1<sup>st</sup> set of moving lines, measured in % movement |
| HL2M | Habituation to 2<sup>nd</sup> set of moving lines, measured in % movement |
| HMSc | Habituation to moth stimulus, measured in % scoot |
| HL1Sc | Habituation to 1<sup>st</sup> set of moving lines, measured in % scoot |
| HL2Sc | Habituation to 2<sup>nd</sup> set of moving lines, measured in % scoot |
| HMB | Habituation to moth stimulus, measured in % burst |
| HL1B | Habituation to 1<sup>st</sup> set of moving lines, measured in % burst |
| HL2B | Habituation to 2<sup>nd</sup> set of moving lines, measured in % burst |
| HME | Habituation to moth stimulus, measured in % escape |
| HL1E | Habituation to 1<sup>st</sup> set of moving lines, measured in % escape |
| HL2E | Habituation to 2<sup>nd</sup> set of moving lines, measured in % escape |
| HMSp | Habituation to moth stimulus, measured in speed (pixels) |
| HL1Sp | Habituation to 1<sup>st</sup> set of moving lines, measured in speed (pixels) |
| HL2Sp | Habituation to 2<sup>nd</sup> set of moving lines, measured in speed (pixels) |
| Home | % time the mouse is inside the red hut |
| Peek | % time the mouse is inside the red hut with its head outside the hut |
| Window | % time the mouse is within 40 pixels of the window |
| Wall | Closest distance from any cage wall (pixels) |
| Food | Closest distance from any food pellet (pixels) |
| DWindow | Distance from the window (pixels) |
| Window Partner | % time spent near the window when partner mouse is also near the window |
| Speed: pixels moved. Movement: speed > 1.5px. Scoot: 11px > speed > 1.5px. Burst: speed > 10px. Escape: speed > 30px. |  |

None of these behavioral parameters showed consistent significant differences between WT and 3xTg-AD mice across all timepoints. Only one behavioral measure (i.e. movement during the first hour of nighttime) showed significant differences in two separate age stages: young (*p* = 0.021), and old (*p* = 0.034), but not aged (*p* = 0.958) mice.

### Unsupervised behavioral classification using Keypoint MoSeq

While both Fiji and DLC are supervised pipelines for behavioral quantification, we also tried an unsupervised behavioral classification approach on the same data. Pose estimation data from DLC featuring 8 labeled mouse body parts was extracted and used to train a Keypoint MoSeq model to recognize discrete behavioral states. These short snippets of behavior - termed “syllables” - capture a wide range of behaviors present across the whole behavioral assay and can be combined to represent complex behavioral paradigms. We trained the model for 550 iterations and reached an average syllable length of 2-4 seconds. We then generated short videos of the mice during each syllable using MoSeq. Visual inspection of these videos was used to manually label each of the 29 syllables (Table 4). These syllables represent a wide array of behaviors that are difficult to quantify using our current supervised methods. We measured the differences in both frequency and duration of each syllable between WT and AD mice at different ages (5-6 month, 12-13 months, and 18-19 months) and found four significantly different syllables: the frequency of syllables 14 and 50 (*p* = 0.025 and *p* = 0.040, respectively), and the duration of syllables 21 and 24 (*p* = 0.018 and *p* = 0.023, respectively). Additionally, we plotted syllable frequency distribution and transition graphs to visualize the differences in genotype across timepoints (Supplementary Figure 1A-F). These graphs suggest that, while there is not a set group of syllables that consistently discriminates between genotypes across aging stages, the MoSeq model is most sensitive to behavioral differences present at 12-13 months of age.

**Table 4.** Top behavioral syllables identified using Keypoint MoSeq.

| Syllable ID | Label | Description |
| --- | --- | --- |
| 0 | groom_hunch | Hunched grooming |
| 1 | burst_turn | Turn 180 and burst across cage length |
| 2 | hunch_look | Hunch, pause, and look in one direction |
| 3 | burst_short | Sudden, agitated, fast burst(s) in a linear direction |
| 4 | freeze_look | Freeze and look around |
| 5 | rear | Rear and look up, while staying in place |
| 6 | stretch_burst | Stretch and take one long step |
| 7 | groom_quick | Grooming in quick separate bursts while looking around |
| 8 | burst_home | Burst and go towards the hut |
| 9 | groom_small | Slow grooming |
| 10 | rear_climb | Rear and/or climb an object or wall |
| 11 | stretch_sit | Stretch upper body forward while sitting and staying in place |
| 12 | ball | Hunched, moving minimally |
| 13 | curl | Stop moving and curl |
| 14 | rear_fast | Rapid rearing around the cage |
| 15 | burst_single | Quick burst away after staying in the same spot |
| 16 | stay | Stay in the same location with minimal movement, not hunched |
| 20 | food | Interact with the food or food plate (i.e. Grab pellet) |
| 21 | stretch | Stretch in place while standing or right after walking |
| 24 | groom_tail | Grooming the tail |
| 27 | loop | Burst away and then circle back to the same spot |
| 28 | sleep | Curled and asleep |
| 31 | groom_still | Grooming while staying still |
| 34 | turn_behind | Turn and/or look behind with a quick posterior groom |
| 42 | turn | Turn while staying in place |
| 46 | look | Look in different directions |
| 50 | turn_away | Turn away and move in opposite direction |
| 61 | freeze_hunch | Stop moving and stay hunched |
| 63 | small_burst | Small movements |

The frequency and duration of each syllable was evaluated across the 22-hour homecage assay. However, more detailed changes in behavior can be captured by measuring syllable frequency and duration throughout key periods of the assay. For instance, focusing on the first 10 minutes after presentation of any stimuli (i.e. moth or moving lines in home quadrant) reveals significant differences in syllable frequency between old WT and 3xTg-AD mice (Supplementary Figure 1G-I). Interestingly, when we looked at period 73 (first presentation of stimuli) we noticed that old 3xTg-AD mice showed a significantly lower frequency of many syllables, exhibiting only 16 out of 29 syllables. This trend was not present on the onset of other stimuli (i.e. periods 85 and 97).

### Therapeutic effect of cyclosporine and previously identified cyclosporine-like drugs on 3xTg-AD mice

We investigated the effect of chronic cyclosporine A, nebivolol, and cabozantinib treatment in young (3 months old) triple-transgenic AD mice (Figure 3A). A dose of 20 mg/kg/day of cyclosporine and 3 mg/kg/day of either nebivolol or cabozantinib was administered in chow through a specially formulated diet (Envigo/Inotiv). Non-treated AD and WT control mice were fed with a base chow diet (2014 Teklad 14%). At 8 months of age, mice on the nebivolol diet were adjusted to a lower dose (0.5 mg/kg/day) due to complications with survivability (Figure 3B).

**Figure 3.**
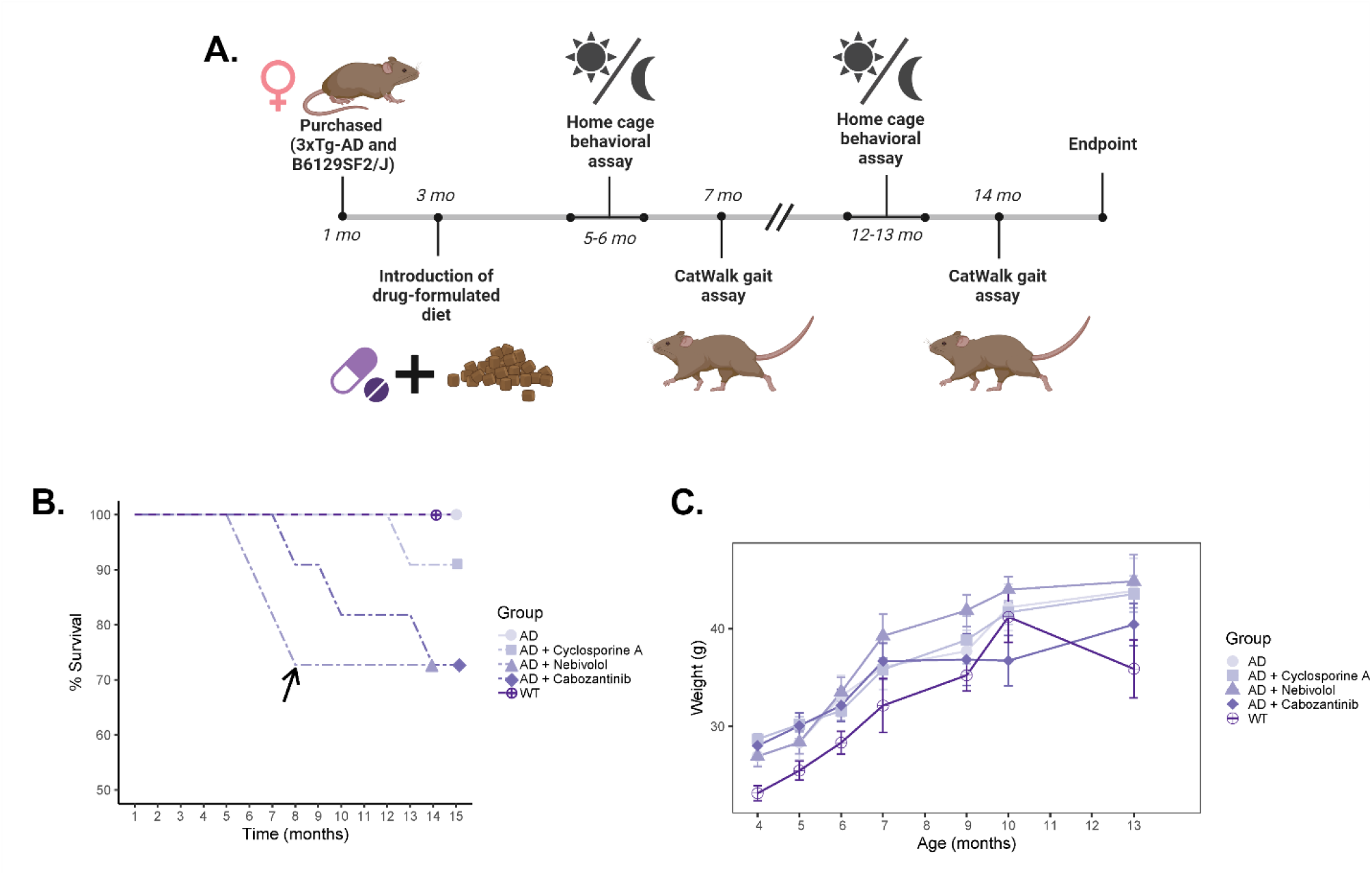
Evaluating the effects of chronic treatment with cyclosporine, nebivolol, or cabozantinib in 3xTg-AD mice. A) Experimental timeline of treatment and behavioral phenotyping in drug-treated 3xTg-AD mice. B) Percent survival of cohorts across experimental time points. The black arrow indicates a change in the administered dose of nebivolol due to decreased survivability. C) Weight measurements across experimental timepoints. Error bars represent SEM. Statistics: two-way ANOVA followed by post hoc Tukey HSD test, * p < 0.05, ** p < 0.01, *** p < 0.001. N = 8 WT mice, N = 16-18 3xTg-AD mice, N = 8-10 3xTg-AD mice treated with cyclosporine, N = 8-10 3xTg-AD mice treated with nebivolol, N = 8-10 3xTg-AD mice treated with cabozantinib.

Using previously determined parameters that revealed gait abnormalities in 3xTg-Ad mice, we examined the effects of these drugs on mice after 4 months of treatment (young) and after 11 months of treatment (old) (Figure 1) (Table 5). Cyclosporine-treated mice showed no significant differences in the number of steps or print length compared to WT mice. The percentage of NSSP was significantly decreased in young cyclosporine-treated mice compared to young WT mice (*p* = 0.029), while the paw angle body axis was significantly increased in the LF paw (*p* = 0.036) and the RH paw (*p* = 0.042). These results suggest a partial rescue of the early gait abnormalities found in 3xTg-AD mice. Additionally, previously observed late gait variations measured using the mean and minimum intensity of the front paws were not present in cyclosporine-treated mice. Moreover, we observed a complete rescue in parameters measuring hind limb support. Nebivolol-treated mice exhibited the same early gait abnormalities as untreated 3xTg-AD mice, but a complete rescue of both late gait variations and hind limb support. Cabozantinib-treated mice showed a complete rescue of early and late gait variations. Interestingly, hind limb support parameters were not significantly different from WT mice, but young cabozantinib-treated mice showed an increase in weight bearing in three of the paws as measured by minimum intensity (LF: *p* = 0.025, RF: *p* = 0.010, LH: *p* = 0.040), as well as both stand (LF: *p* = 0.022, RF: *p* = 0.008, LH: *p* = 0.007, RH: *p* = 0.019) and step cycle (LF: *p* = 0.024, RF: *p* = 0.008, LH: *p* = 0.021, RH: *p* = 0.036) in all four paws. These differences were not significant in older mice. Overall, we observed varying levels of rescue from drug treatments, with cyclosporine and nebivolol being the most effective at preventing early gait abnormalities, and all treatments rescuing late gait abnormalities.

**Table 5.**
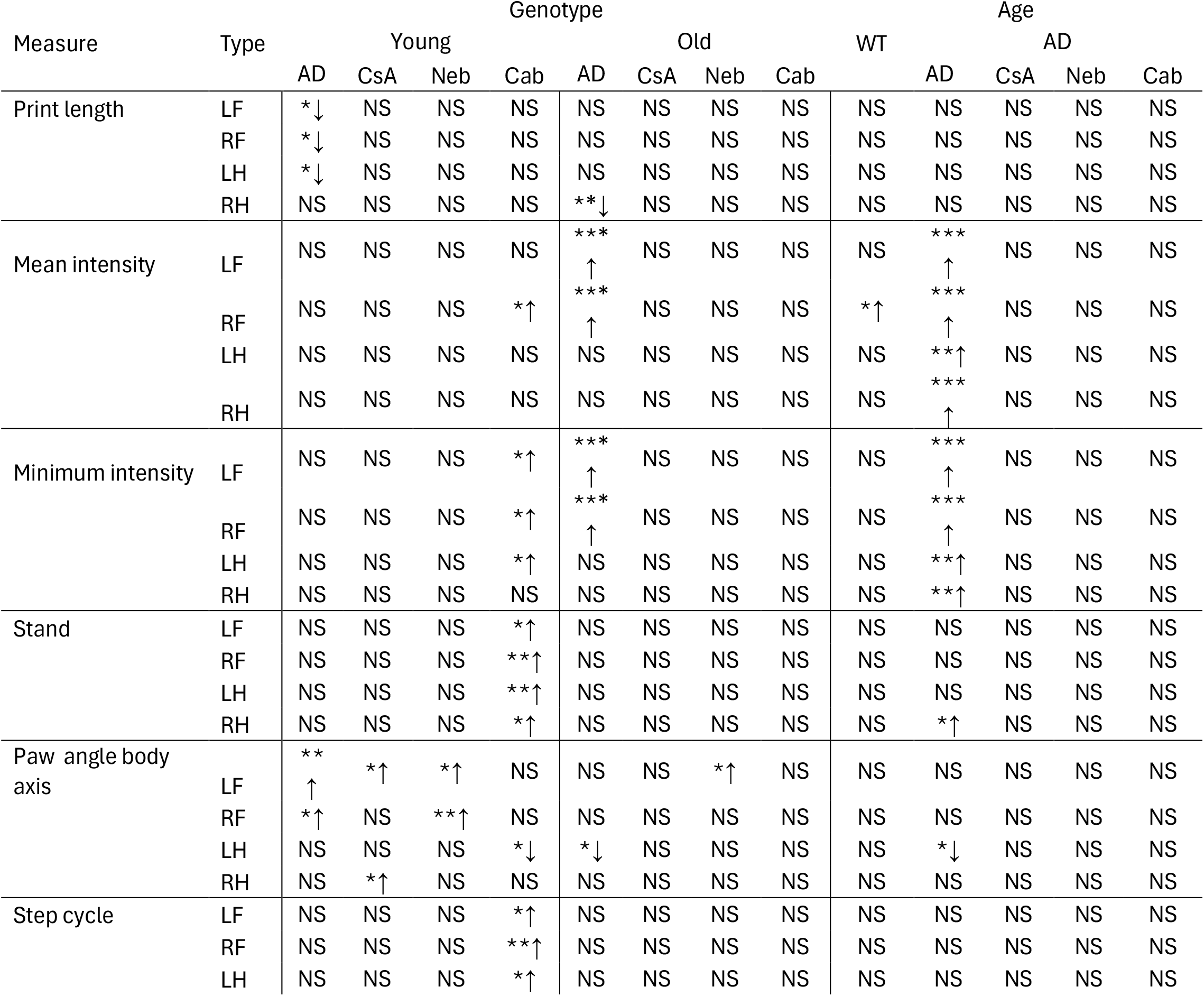
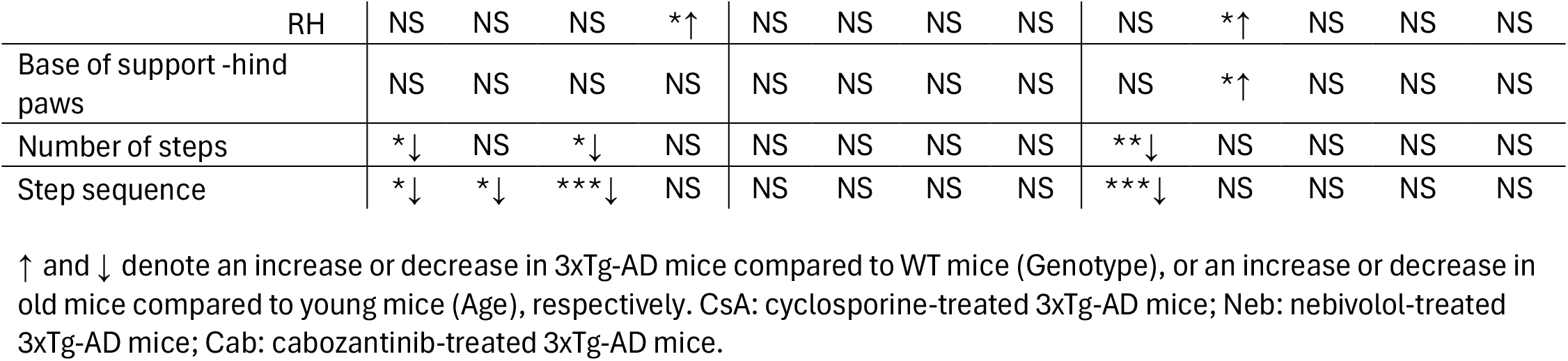
CatWalk gait analysis of drug-treated 3xTg-AD mice.

Mice were tested in our custom built 8-cage imaging system at 5-6 months of age (young) and again at 12-13 months of age (old). Through the use of animal tracking, pose estimation, and unsupervised behavioral classification, we were able to obtain a large number of behavioral parameters measuring a wide range of behaviors. Notably, we captured behaviors defined by both simple measures (e.g. activity, time spent in cage quadrants) and complex measures (e.g. sociability, time spent eating, grooming, or peeking from inside the hut) in an automated fashion. Thus far, we have analyzed these parameters independently to both validate and identify novel behavioral markers in 3xTg-AD mice. While these scalar behavioral measures are useful for understanding independent trends found within a disease model, they manifest in an age-dependent manner that is not appropriate for discriminating between disease states, especially in longitudinal studies. Therefore, we combined the behavioral parameters generated by each analysis method (Fiji, DeepLabCut, MoSeq) to produce 125-parameter behavioral profiles for all the experimental groups (Supplementary File 3).

We performed principal component analysis (PCA) to assess the underlying patterns within the behavioral data (Figure 4A). The first two components accounted for 82% of the variance. PC1 was characterized by high loadings for variables that differentiate between young and old WT mice, while PC2 was primarily associated with variables that separate across the genotypes. We then calculated weights for each of the 125 behavioral parameters using the loadings of each variable from PC1. This weighted data was used to perform hierarchical cluster analysis to explore the grouping of drug-treated 3xTg-AD mice with untreated and WT controls (Figure 4B). The resulting dendrogram features three main clusters: a cluster of old WT mice with cyclosporine-treated mice and young nebivolol-treated mice, a cluster of young WT and 3xTg-AD with old drug-treated mice, and a cluster of old 3xTg-AD mice with young cabozantinib-treated mice. Visual inspection of these three clusters reveals two oppositely distinct sets of behavioral profiles: the old WT cluster and the old 3xTg-AD cluster. However, the cluster containing both young WT and 3xTg-AD mice is characterized by an intermediate behavioral profile without a clear pattern. These clustering results further show a high sensitivity to differences between old, but not young, WT and 3xTg-AD mice.

**Figure 4.**
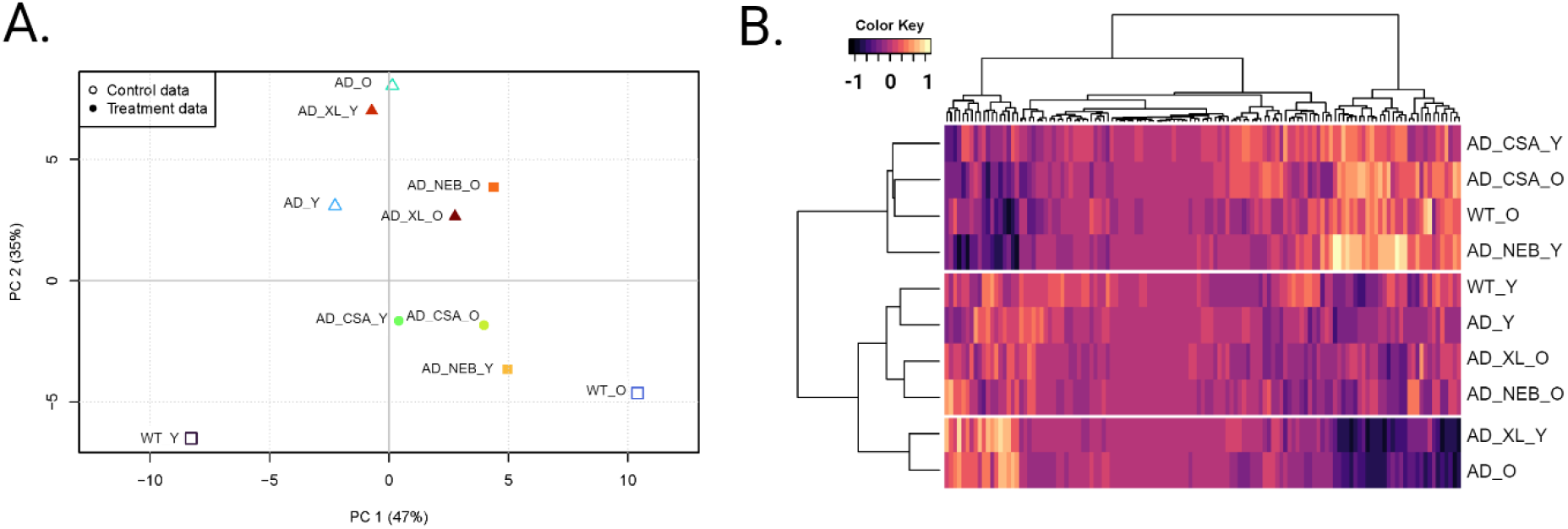
Cluster analysis. A) Principal component analysis (PCA) and B) hierarchical cluster analysis of the 125-parameter behavioral profiles. WT_Y: young wild-type mice; WT_O: old wild-type mice; AD_Y: young 3xTg-AD mice; AD_O: old 3xTg-AD mice; AD_CSA_Y: young cyclosporine-treated 3xTg-AD mice; AD_CSA_O: old cyclosporine-treated 3xTg-AD mice; AD_NEB_Y: young nebivolol-treated 3xTg-AD mice; AD_NEB_O: old nebivolol-treated 3xTg-AD mice; AD_XL_Y: young cabozantinib-treated 3xTg-AD mice; AD_XL_O: old cabozantinib-treated 3xTg-AD mice.

Based on these analyses, we conclude that the cyclosporine, nebivolol, and cabozantinib treatments have beneficial effects. Old 3xTg-AD mice treated with cyclosporine display a similar behavioral profile as old WT mice, indicating that cyclosporine suppresses the effects of AD risk genes in the 3xTg-AD line. Similarly, old AD-model mice treated with nebivolol and cabozantinib display a behavioral profile similar to young 3xTg-AD mice, suggesting that these drugs prevent age-associated effects in this triple-transgenic AD line.

## Discussion

The triple transgenic mouse model of Alzheimer’s disease (3xTg-AD) is widely used to study the progression of the disease in multiple contexts, including motor function^17^. While physical changes are not as pronounced as the decline of cognitive function, they present more consistently during the mild to moderate stages of the disease^18^. In the present study, we evaluated changes in the motor function and gait of 3xTg-AD mice during the early and late stages of the disease. Our results point toward differences in the positioning of the paws as well as the sequence of steps and patterns, which become more subtle as the mice age. We also found a higher tendency for 3xTg-AD to use their hind limbs for support as they age. These results are consistent with those found in other models of Alzheimer’s disease^19^. Mice treatment with cyclosporine, nebivolol, or cabozantinib showed partial rescue of these gait abnormalities – during both early and late stages of treatment.

Classic behavioral assays often place the animal in “unnatural” environments in order to test habituation, short- and long-term memory, or associative learning. In mouse models, batteries of tests often include the Morris Water Maze, Open Field test, and Novel Object Recognition, all of which involve multiple short exposures to standard testing chambers. While these tests are able to capture important behavioral phenotypes that are relevant to the disease, they are often subject to interstudy variability. This is especially highlighted in the context of rodent Alzheimer’s disease (AD) models, where differences in model, age, sex, and cohorts can alter study results^10,20-22^. Automated analysis of behavior is a key tool for spotting subtle differences in behavioral phenotypes in a minimally biased manner. In this study, we have compiled multiple freely available and open-source tools to enhance the behavioral analysis from videos obtained in a low-cost timelapse setup. This high-throughput pipeline is capable of imaging 8 mice at once for 22 consecutive hours, requiring a low amount of human intervention while still capturing a wide range of behavioral measures. The resulting behavioral profiles generated from these measures are capable of differentiating between both age and genotype.

We were able to validate the effectiveness of chronic cyclosporine treatment in a triple-transgenic mouse model of AD, during both early and late stages of the disease. Additionally, we looked at the effects of chronic nebivolol and cabozantinib treatment in these mice. Nebivolol is an FDA-approved β1 adrenergic receptor antagonist used in the treatment of hypertension. Previous studies have shown promising effects in both *in vitro* and *in vivo* models of AD^23,24^. Links between hypertension and AD point to the relevance of cardiovascular factors in the progress of the disease^25-27^. Cabozantinib is a tyrosine kinase inhibitor which targets vascular endothelial growth factor receptor 2 (VEGFR2). It is often prescribed to treat medullary thyroid cancer and some kidney and liver cancers. A recent study found that cabozantinib partially reduced tau hyperphosphorylation and astrogliosis in 5xFAD mice^28^. Protein kinase inhibitors have been previously proposed as candidates for AD treatment due to their autophagy modulation properties^29^. Treatment with nebivolol and cabozantinib resulted in partially ameliorating effects of AD behavioral phenotypes. It is likely that a dose adjustment for both drugs is required to achieve optimal preventative effects in this model. Future studies should focus on optimizing drug dosage and delivery methods to achieve optimal therapeutic effects, as well as evaluating the underlying molecular mechanisms by which these candidate drugs ameliorate the behavioral pathology of AD models.

## Methods

### Ethics declaration and approval of animal experiments

The present study was performed in compliance with federal regulations and guidelines regarding the ethical and humane use of animals, and followed PREPARE, 3R, and ARRIVE 2.0 guidelines to plan and carry out the experiments. All protocols and experimental procedures were approved by Brown University’s Institutional Animal Care and Use Committee (Animal Welfare Assurance Number D16-00183).

### Animals

In this study, we used 3xTg-AD (JAX stock #004807) mutant mice (n = 50), and nontransgenic B6129SF2/J (JAX stock #101045) wild-type control mice (n= 20) obtained from the Jackson Laboratory (Bar Harbor, ME, USA). Groups of 4-6 mice were housed in Optimice IVC cages located in Brown University’s animal care facility with *ad libitum* access to diet and water. The animal room was maintained on a 12-hour light/dark cycle. Starting at 3 months of age, 3xTg-AD mice were given either a standard mouse diet (2014 Teklad Global 14% protein diet) or a formulated mouse diet containing either Cyclosporine A (150 ppm), Nebivolol (23 ppm), or Cabozantinib (23 ppm) purchased from Envigo/Inotiv (Madison, WI, USA). The drugs selected for this study were chosen based on our previous screening for CsA-like FDA-approved drugs in a zebrafish larvae model^5^. The administered concentration of each drug was adapted from 10µM based on previous literature in mouse models^23,30,31^. Mice were routinely weighed to ensure proper consumption of the newly introduced diets and to check overall health during chronic drug treatments.

### Analysis of gait function

Gait recording and analysis was conducted in the Carney Institute Rodent Behavioral Phenotyping Facility at Brown University. We used a semi-automated gait analysis method (CatWalk XT 10.6; Noldus Information Technology Inc.) to obtain and quantify high-resolution gait function data^32,33^. The CatWalk system is capable of obtaining several mobility measures through a backlit camera illuminating a glass stage where the mice are allowed to freely walk from one end to another. We measured multiple temporal and spatial gait parameters, including speed, step patterns, and cadence (Table 1). Mice were transported to the experimental room with the CatWalk apparatus and allowed to stay in their home cages for 30 minutes. Each mouse was then weighed and placed in a separate cage before being moved to the edge of the glass walkway to begin the experiment. Acquisition started as soon as the mouse was placed on the walkway and stopped once the mouse successfully walked three times across the walkway, at which point the mouse was returned to its home cage. A compliant run was categorized as a one-directional walk from one end of the recording area to another performed in less than 10 seconds. All non-compliant runs were excluded from further analysis. Before the start of each experimental session, a representative control mouse was used to determine optimal settings to correctly detect paw prints and separate them from background noise. Food pellets were placed on the goal box to encourage mice to traverse the walkway. We used 70% ethanol to thoroughly clean the apparatus between mice.

### The 8-cage imaging system

Time lapse recordings of n=60 mice in our 8-cage system were obtained as previously described^16^. Briefly, for each experimental session, 8 mice were placed individually in white polypropylene cage bottoms (Thomas Scientific, Cat No. 1113H35) with access to food and water. All mice had visual access to a non-littermate mouse through a 2.5-inch diameter hole located in the lower inside-facing corner of each cage (Figure 2A). A red hut was also placed inside each cage for shelter (Bio-Serv, K3357). We recorded all experiments using the combined video stream of two webcams (Logitech C922x Pro) at a snapshot interval of 1s, total image size of 640 x 960 pixels, and using the ffdshow video codec for video compression in the open-source software SkyStudioPro (version 1.1.0.30, skystudiopro.com).

The behavioral assay consists of a 22-hour PowerPoint presentation simulating a daytime period (6 hours, white background), a nighttime period (6 hours, red background), a 6-hour nighttime period with stimuli presented in the form of a rotating yellow moth and moving red lines, and a 4-hour daytime period (Figure 2B, Supplementary File 1). The day/night transitions were timed to match the mice’s standard light/dark cycle in the animal housing room. All behavioral assays were started at 2pm and ended at 12pm on the following day. Cages were cleaned with 70% ethanol at the end of each experiment.

### Generation of behavioral profiles

Fiji: Automated tracking and analysis of the 8-cage videos was performed in Fiji using a previously developed macro (Supplementary File 2)^16^. The macro allows the user to track the mouse centroid and head position during both daytime and nighttime by separating the color channels and utilizing the red channel for subsequent analyses. We generated a total of 15 behavioral parameters measuring movement, acclimation, habituation, and location. A list of all parameters and their descriptions can be found in Table 2.

DeepLabCut: In addition to Fiji, we also used our previously published DeepLabCut 2.2.3 (DLC) model to perform pose estimation in our 8-cage videos^16^. We trained the network (based on ResNet-50) to recognize 12 mouse body parts,15 cage features, and 21 different stimuli (Supplementary figure 2, https://github.com/Creton-Lab). Videos were cropped to outline each of the 8 cages using a custom Python script; a total of 430 frames from 10 videos were used to train the network for 400,000 iterations. The output files generated after running DLC on the video files (.csv files containing x,y coordinates and a likelihood percentage for each body part) were processed using customized Python scripts to filter for low likelihood measurements (< 0.90) and to calculate 52 behavioral parameters encompassing activity, sociability, and habituation to stimuli. A list of each parameter - along with descriptions - is outlined in Table 3.

Keypoint MoSeq: Positional features extracted from DLC were used as input to train a MoSeq model - an unsupervised machine learning method for behavioral classification^34^. We used 8 labeled mouse body parts (nose, right ear, left ear, head, neck, center, hip, tail base) as input for Keypoint Moseq; these features are similar to the default recommended labeling scheme. In brief, MoSeq uses unsupervised machine learning to classify behaviors into motifs or “syllables” that represent behavioral states frequently used in the mice from the input data. These syllables are segments of complex behavioral fingerprints that can be used to discriminate between experimental groups and/or treatments. DLC data from 26 videos was used as input for the Keypoint MoSeq pipeline. The Keypoint MoSeq model was trained for 550 iterations converging on an average syllable length of 2-4 seconds. In order to produce syllables capturing the whole breadth of the experimental groups, we included videos from each genotype and treatment for the model training. The trained model produced 29 syllables explaining 90% of the variance in behavior across the home cage behavioral assay.

### Hierarchical clustering and Principal Component Analysis

The behavioral responses of each mouse were summarized to form behavioral profiles for each experimental group: untreated 3xTg-AD mice (AD), 3xTg-AD mice treated with cyclosporine A, nebivolol, or cabozantinib, and B6129SF2/J control mice (WT). A total of 125 behavioral measurements were used to create the behavioral profiles. Due to similarities of some of the behaviors (e.g. speed during the first hour and burst movement during the first hour), we introduced weights for each behavior. Weights were calculated using Principal Component Analysis (PCA) and assigned according to the loading of each measure on principal component 1 (PC1). These measures were then scaled and used for hierarchical cluster analysis using the Euclidean distance similarity metric with Ward’s method. All data visualizations were generated in R (version 4.3.2) and arranged in BioRender (BioRender.com).

### Statistics

Statistical analyses were completed using R (version 4.3.2). Tests for normality were conducted using the Shapiro-Wilk test. For gait analysis, two-way analysis of variance (ANOVA) followed with a Tukey HSD post hoc test when a significant interaction was observed was used for comparisons to controls. Age, genotype, and age-by-genotype interactions were considered. For statistical analysis of the behavioral parameters measured with our homecage assay, we performed the non-parametric unpaired two-samples Wilcoxon test. Kruskal–Wallis and post hoc Dunn’s two-sided test were used to evaluate differences in syllable frequencies generated by Keypoint MoSeq. Significance for all statistical tests was set at p < 0.05.

## Data sharing and availability

Our trained DLC model, as well as all pre- and post-processing Python scripts are available on GitHub (https://github.com/Creton-Lab). All data are available in the main text or the supplementary materials.

## Supplementary Materials

**Supplementary Figure 1**. Analysis of syllables generated with Keypoint MoSeq. Frequency of syllables in A) young, B) old, and C) aged 3xTg-AD mice compared to WT mice. Syllable transition graphs of D) young, E) old, and C) aged 3xTg-AD mice compared to WT mice. Syllable frequencies of old mice during the first 10 minutes of the moth stimulus (G), the first set of moving lines in the home quadrant (H), and the second set of moving lines in the home quadrant (I).

**Supplementary Figure 2**. Mouse, cage, and stimuli markers used in the DeepLabCut model. 27 out of 48 markers were utilized for analysis in the current study.

**Supplementary File 1**. PowerPoint Presentation of 22-hour behavioral assay.

**Supplementary File 2**. Fiji/ImageJ macro for the tracking and analysis of mouse behavior in an 8-cage imaging system.

**Supplementary File 3**. Raw data with measures of behavior in Fiji, DeepLabCut, and MoSeq. Columns d0-d63 = duration of motion sequences (some motion sequences, such as d17-19, are automatically removed by the software). Columns f0-f63 = frequency of motion sequences. Columns M1-Win Partner = behaviors measured in DeepLabCut. Columns M1-Out = behaviors measured in Fiji. Each row represents an individual mouse.

**Supplementary Table 1.** CatWalk gait analysis.

| Measure | Type | Genotype |  | Age |  |
| --- | --- | --- | --- | --- | --- |
|  |  | Young | Old | WT | AD |
| Print length | LF | *↓ | NS | NS | NS |
|  | RF | *↓ | NS | NS | NS |
|  | LH | *↓ | NS | NS | NS |
|  | RH | NS | **↓ | NS | NS |
| Toe spread | LH | NS | NS | NS | NS |
|  | RH | NS | NS | NS | NS |
| Intermediate toe spread | LF | NS | NS | NS | NS |
|  | RF | NS | NS | NS | NS |
|  | LH | NS | NS | ***↓ | ***↓ |
|  | RH | NS | NS | ***↓ | ***↓ |
| Print area | LF | *↑ | NS | *↑ | NS |
|  | RF | NS | NS | *↑ | NS |
|  | LH | NS | NS | NS | NS |
|  | RH | NS | NS | NS | NS |
| Maximum contact area | LF | **↑ | NS | *↑ | NS |
|  | RF | NS | NS | *↑ | NS |
|  | LH | NS | NS | NS | NS |
|  | RH | NS | NS | NS | NS |
| Mean intensity | LF | NS | ***↑ | NS | ***↑ |
|  | RF | NS | ***↑ | *↑ | ***↑ |
|  | LH | NS | NS | NS | **↑ |
|  | RH | NS | NS | NS | ***↑ |
| Minimum intensity | LF | NS | ***↑ | NS | ***↑ |
|  | RF | NS | ***↑ | NS | ***↑ |
|  | LH | NS | NS | NS | **↑ |
|  | RH | NS | NS | NS | **↑ |
| Stand | LF | NS | NS | NS | NS |
|  | RF | NS | NS | NS | NS |
|  | LH | NS | NS | NS | NS |
|  | RH | NS | NS | NS | *↑ |
| Paw angle body axis | LF | **↑ | NS | NS | NS |
|  | RF | *↑ | NS | NS | NS |
|  | LH | NS | *↓ | NS | *↓ |
|  | RH | NS | NS | NS | NS |
| Step cycle | LF | NS | NS | NS | NS |
|  | RF | NS | NS | NS | NS |
|  | LH | NS | NS | NS | NS |
|  | RH | NS | NS | NS | *↑ |
| Support | Lateral | NS | NS | NS | NS |
|  | Girdle | NS | NS | NS | NS |
|  | Diagonal | NS | NS | NS | NS |
|  | Four | NS | NS | NS | NS |
|  | Three | NS | NS | NS | NS |
|  | Single | NS | NS | NS | NS |
|  | Zero | NS | NS | NS | NS |
| Base of support -front paws |  | NS | NS | NS | NS |
| Base of support -hind paws |  | NS | NS | NS | *↑ |
| Duration |  | NS | NS | NS | NS |
| Cadence |  | NS | NS | NS | NS |
| Number of steps |  | *↓ | NS | **↓ | NS |
| Step sequence |  | *↓ | NS | ***↓ | NS |

↑ and ↓ denote an increase or decrease in 3xTg-AD mice compared to WT mice (Genotype), or an increase or decrease in old mice compared to young mice (Age), respectively.

## Acknowledgements

We thank Dr. Arturo Andrade for his valuable input and support with the gait analysis. This work was supported by the National Institutes of Health, Grant R01 GM136906 (R.C.) and Grant R01 GM136906-03S1 (R.C., J.A.K.).

