## Supplementary figures and images for "Combining supervised and unsupervised analyses to quantify behavioral phenotypes and validate therapeutic efficacy in a triple transgenic mouse model of Alzheimer’s disease"

### Supplementary Figure 1

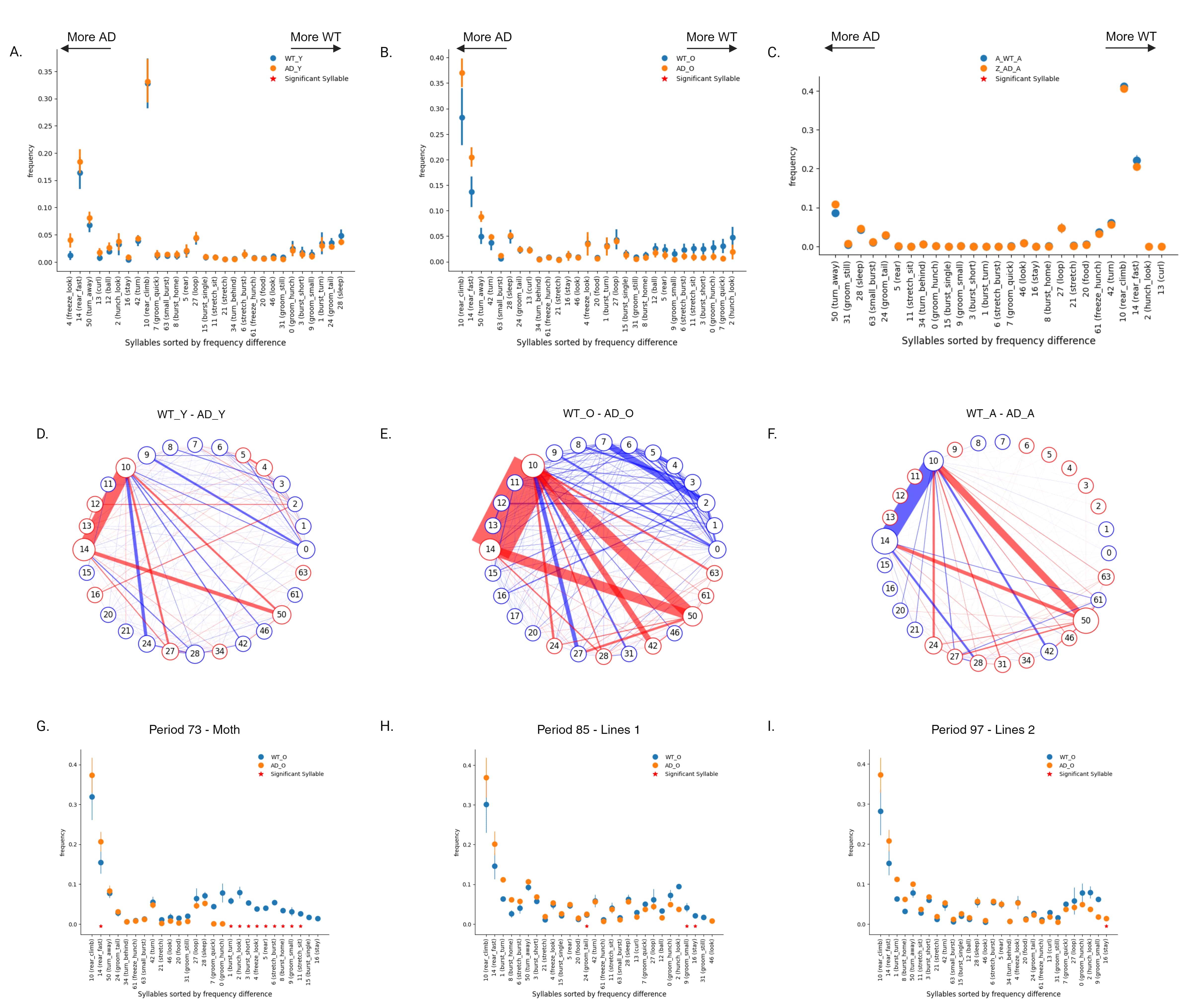

### Supplementary Figure 2

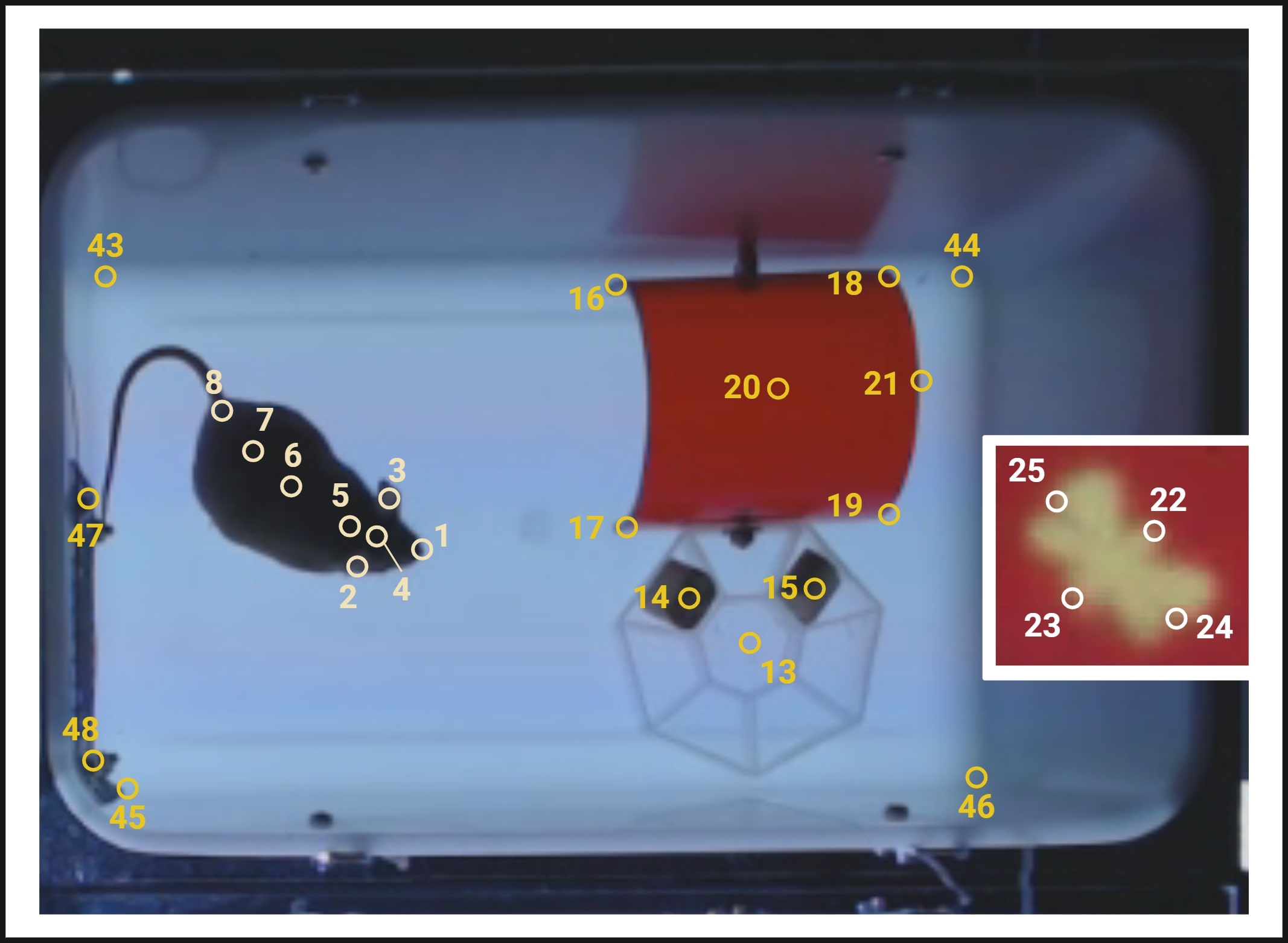
