## Supplementary File 1 for "Combining supervised and unsupervised analyses to quantify behavioral phenotypes and validate therapeutic efficacy in a triple transgenic mouse model of Alzheimer’s disease"

### Slide 1
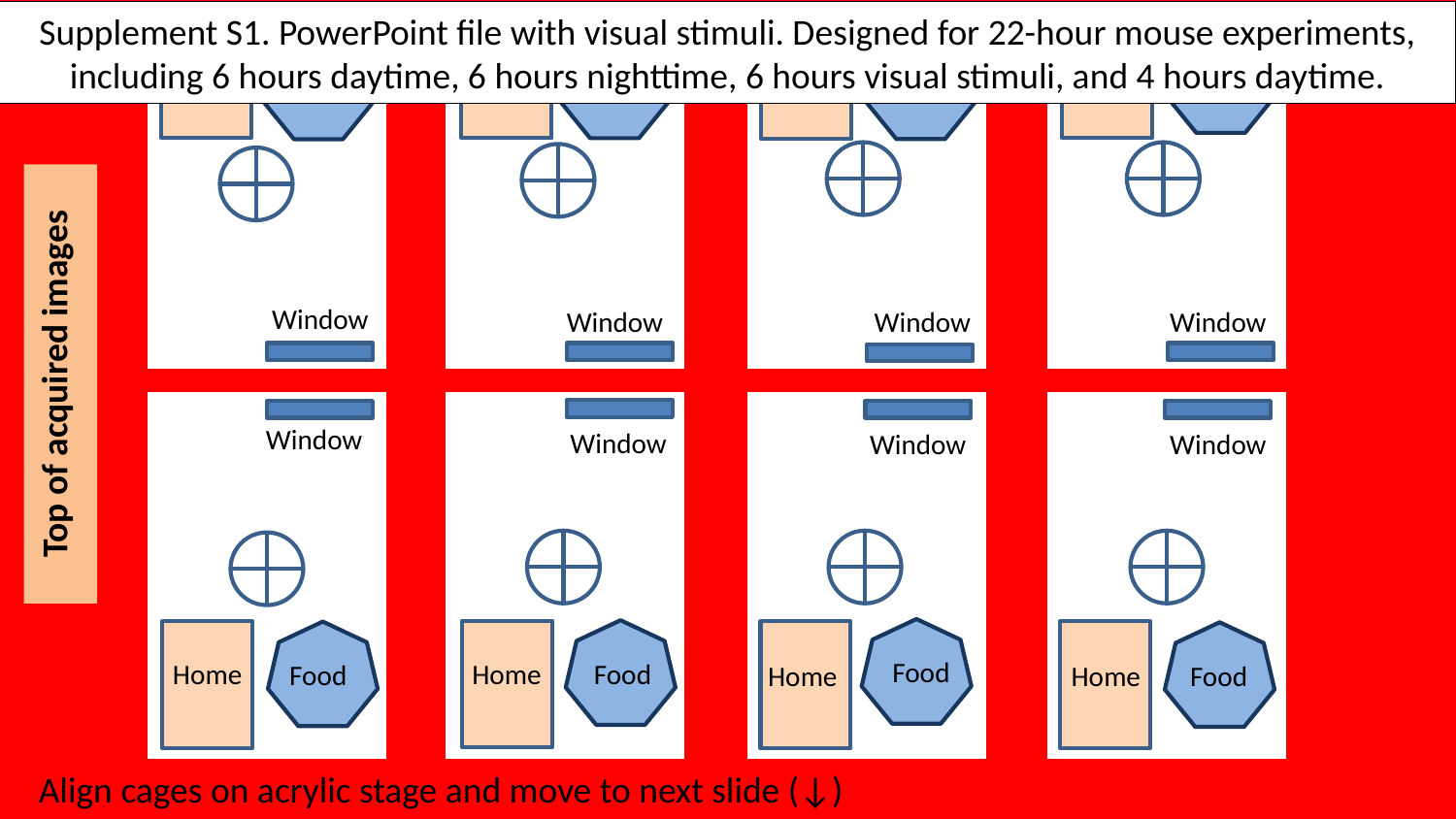

Supplement S1. PowerPoint file with visual stimuli. Designed for 22-hour mouse experiments, including 6 hours daytime, 6 hours nighttime, 6 hours visual stimuli, and 4 hours daytime.
Food
Food
Home
Food
Home
Home
Food
Home
Food
,,
Top of acquired images
Window
Window
Window
Window
Window
Window
Window
Window
Food
Home
Home
Food
Food
Food
Home
Home
Align cages on acrylic stage and move to next slide (↓)

### Slide 2
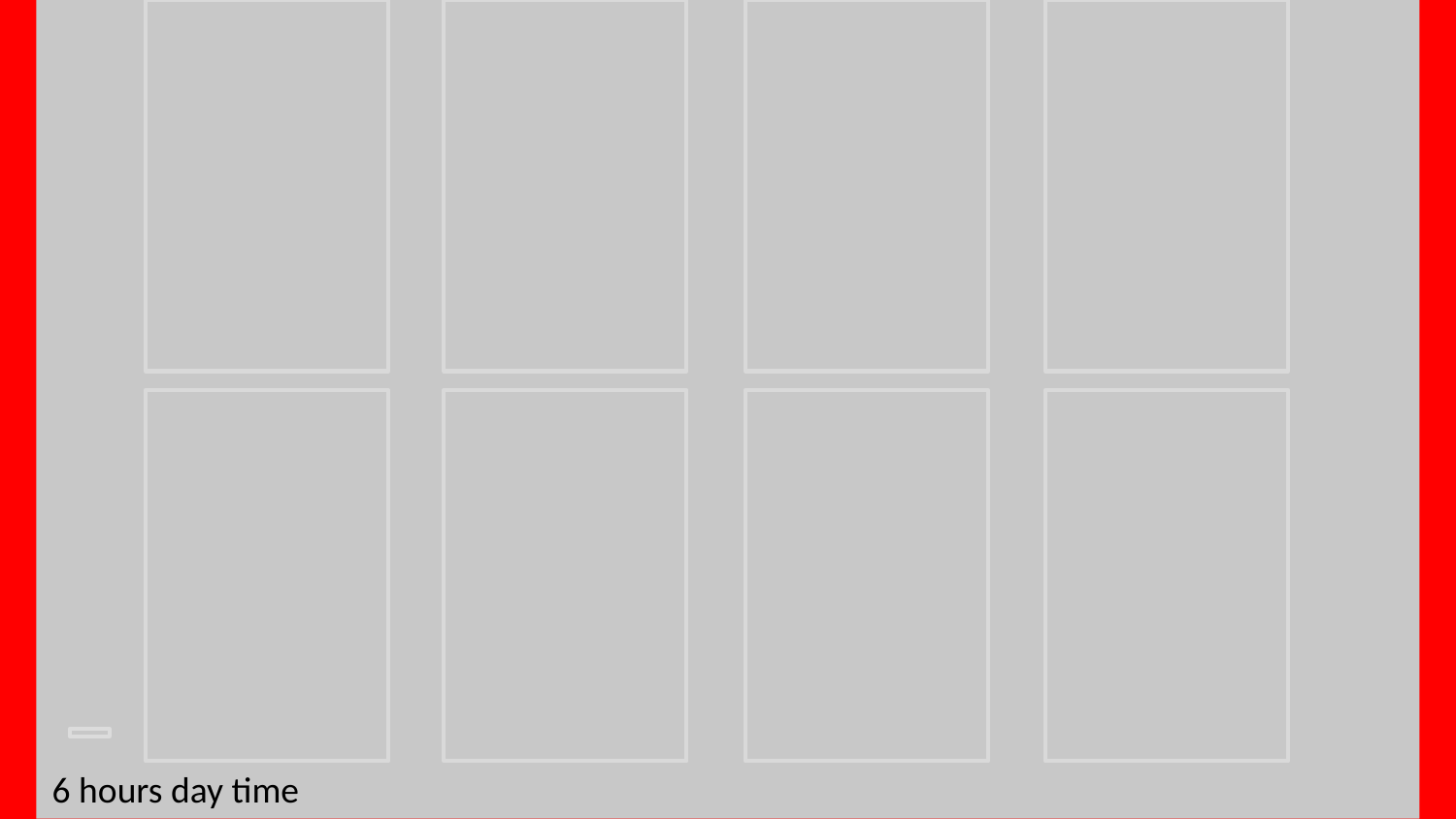

6 hours day time

### Slide 3
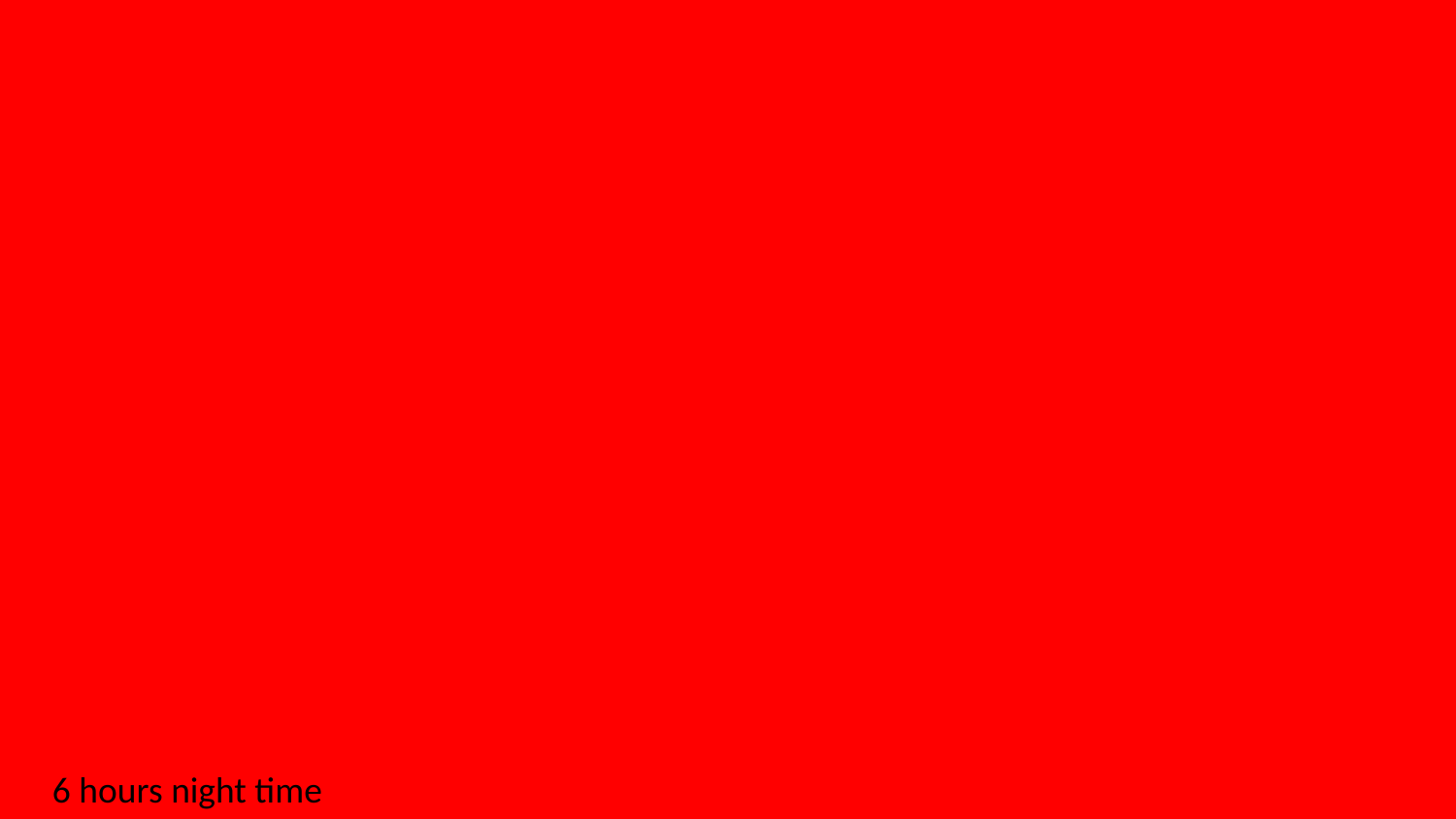

6 hours night time

### Slide 4
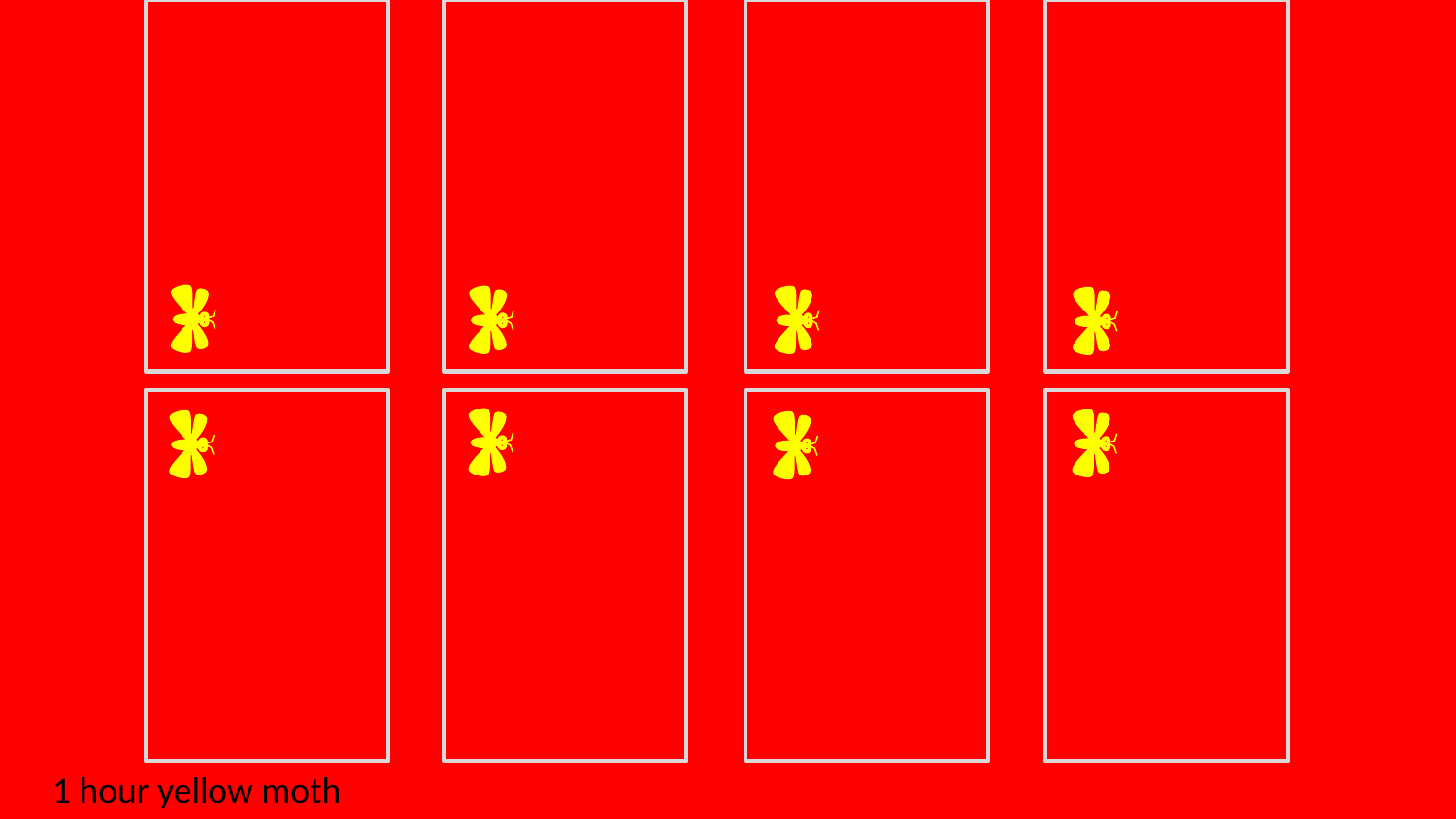

1 hour yellow moth

### Slide 5
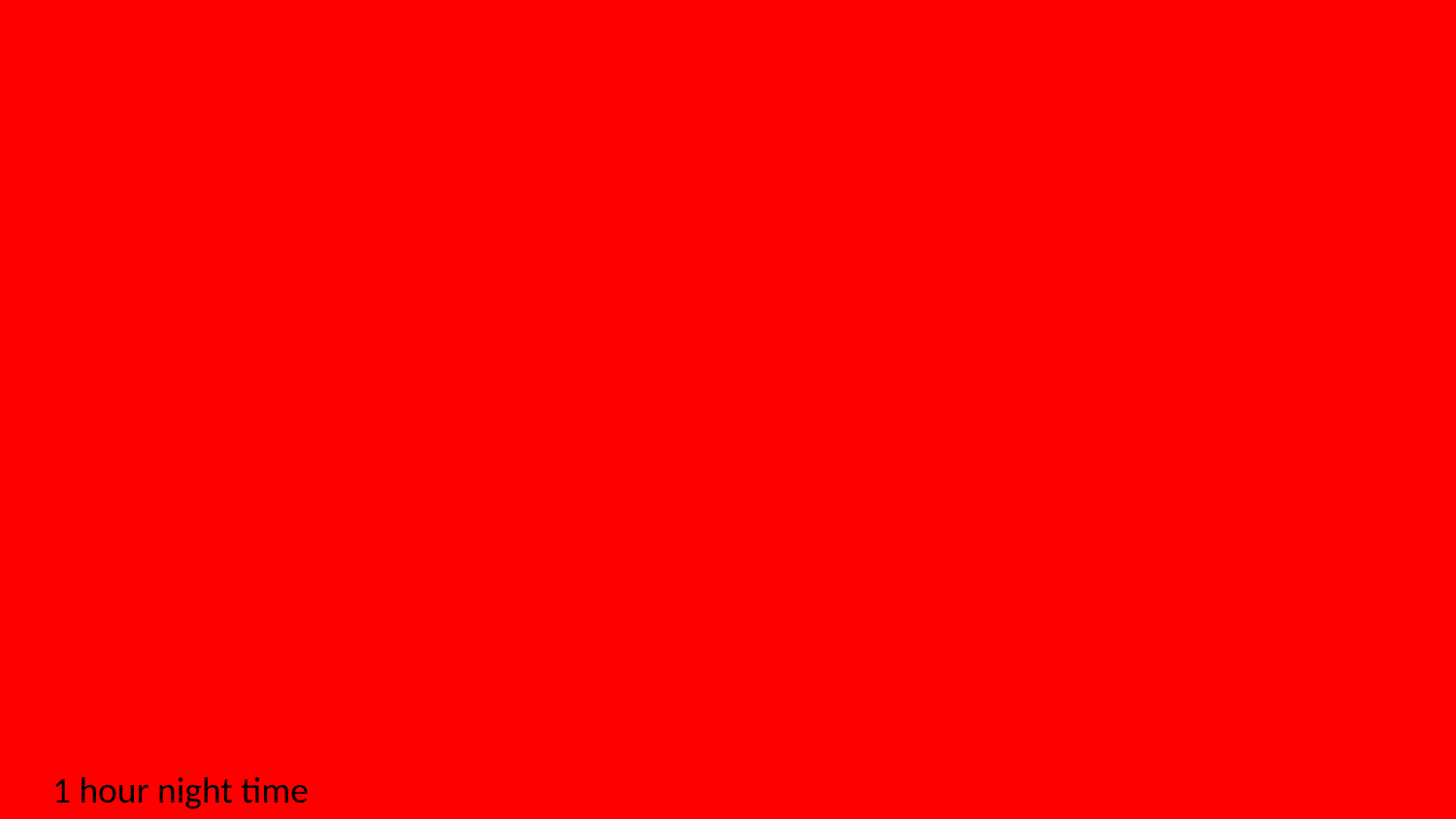

1 hour night time

### Slide 6
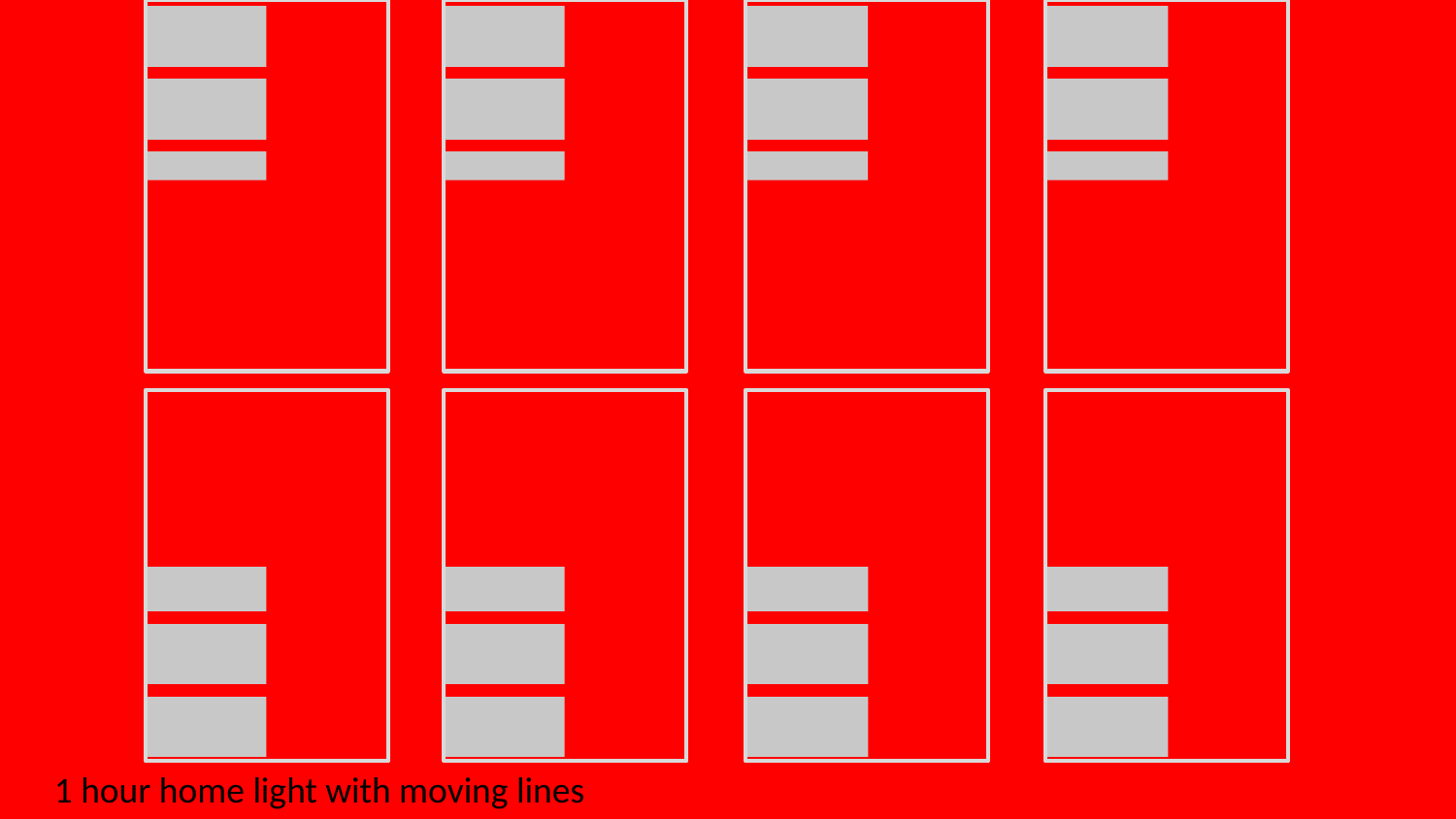

1 hour home light with moving lines

### Slide 7
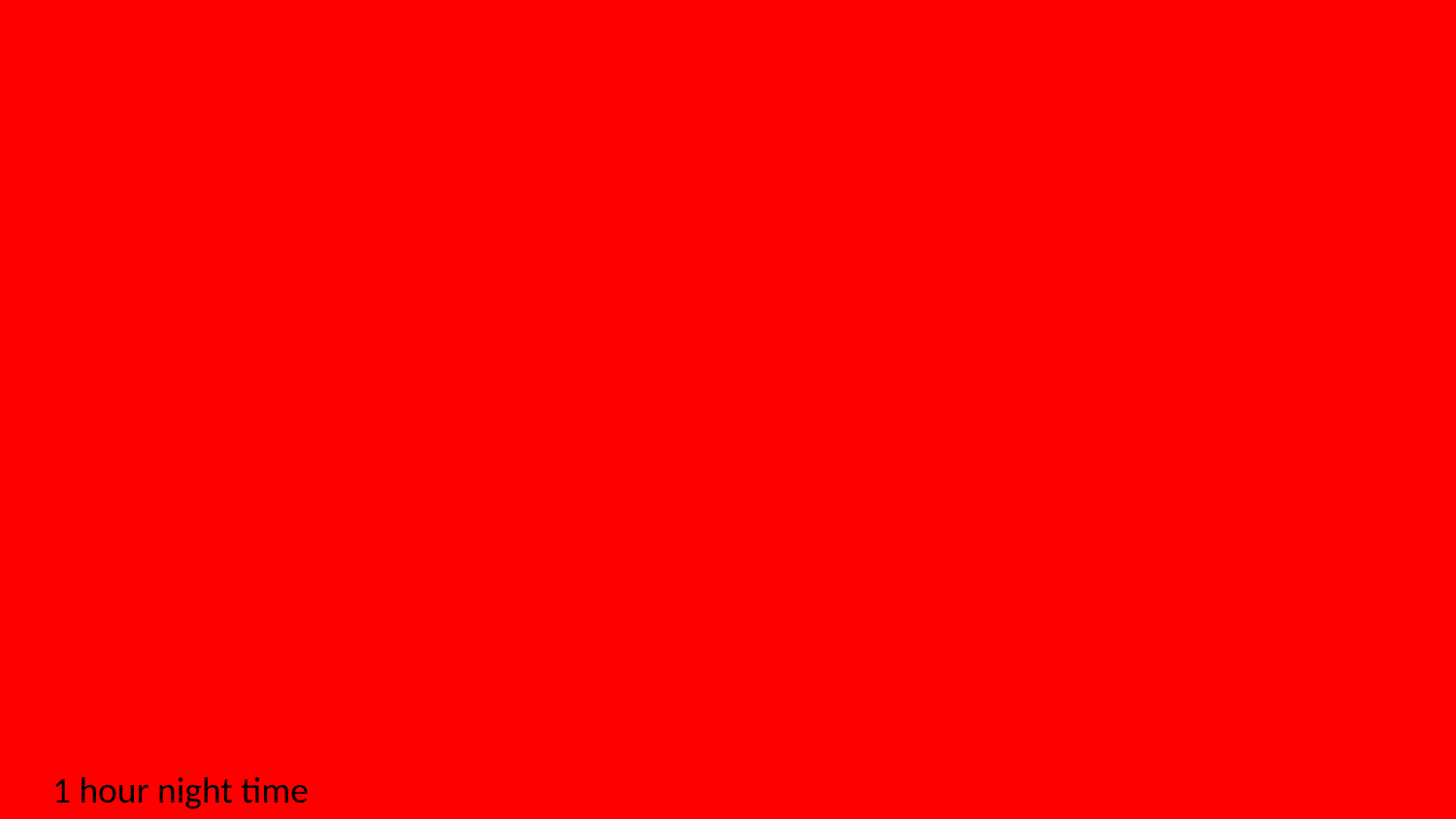

1 hour night time

### Slide 8
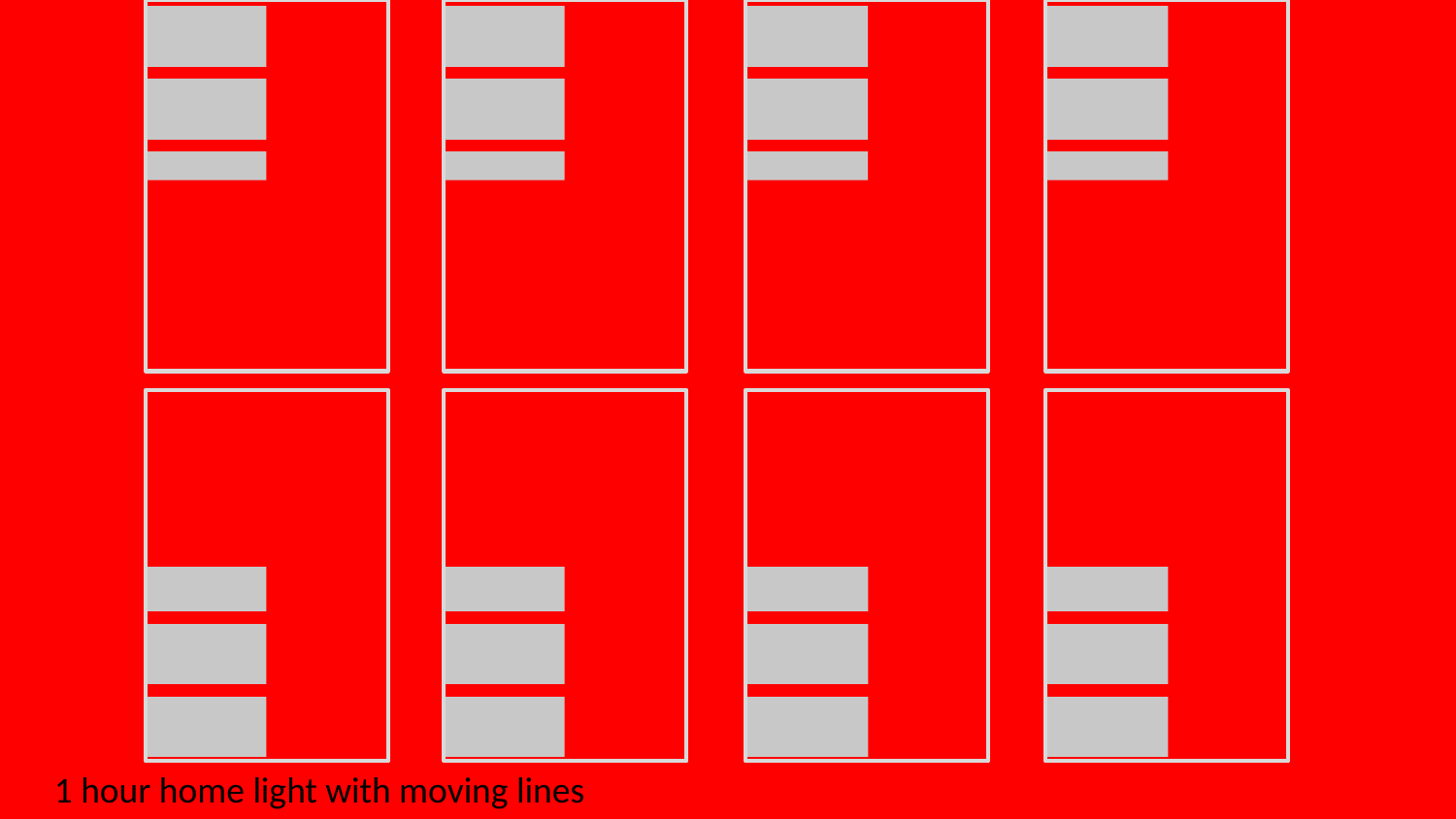

1 hour home light with moving lines

### Slide 9
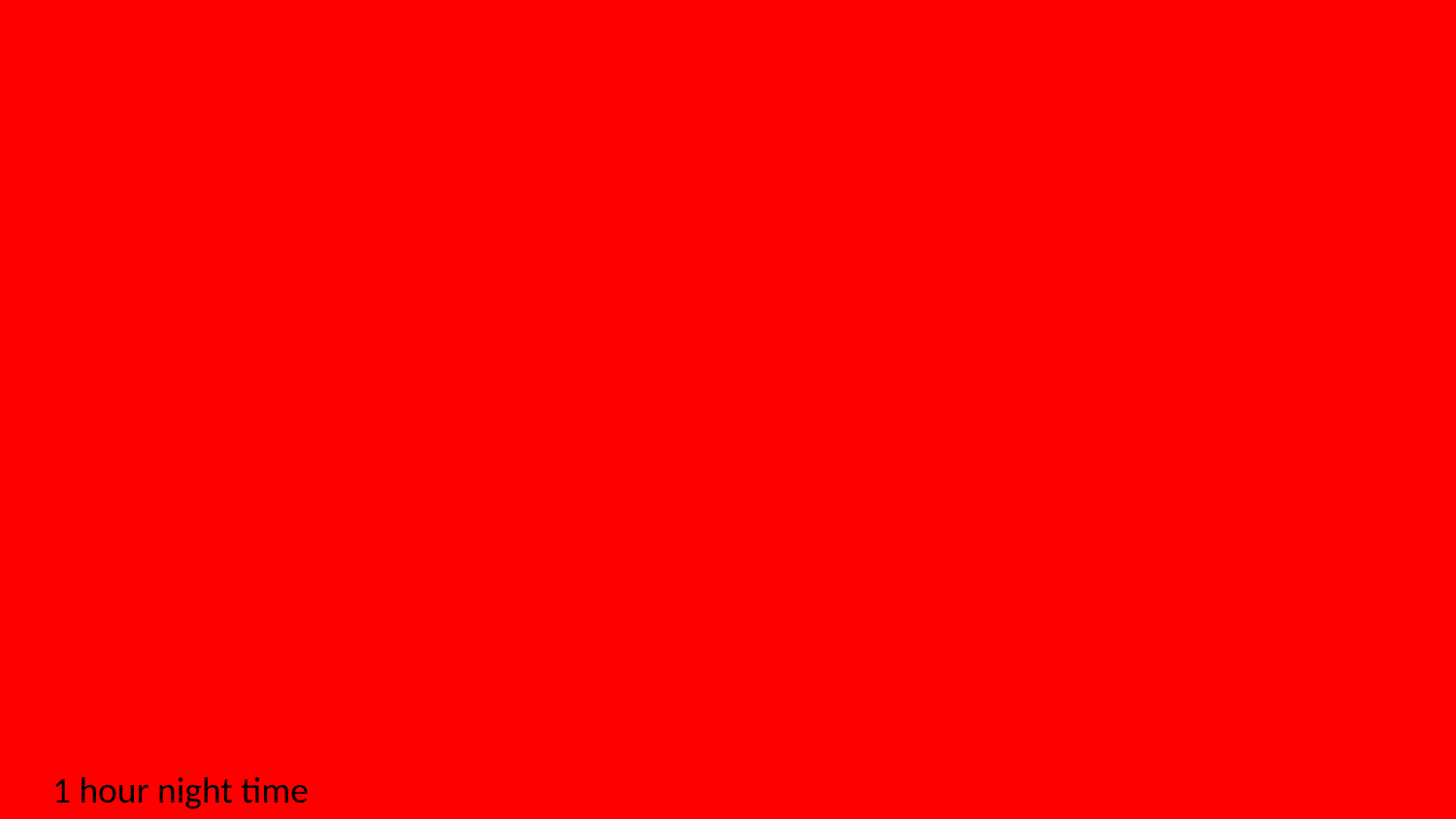

1 hour night time

### Slide 10
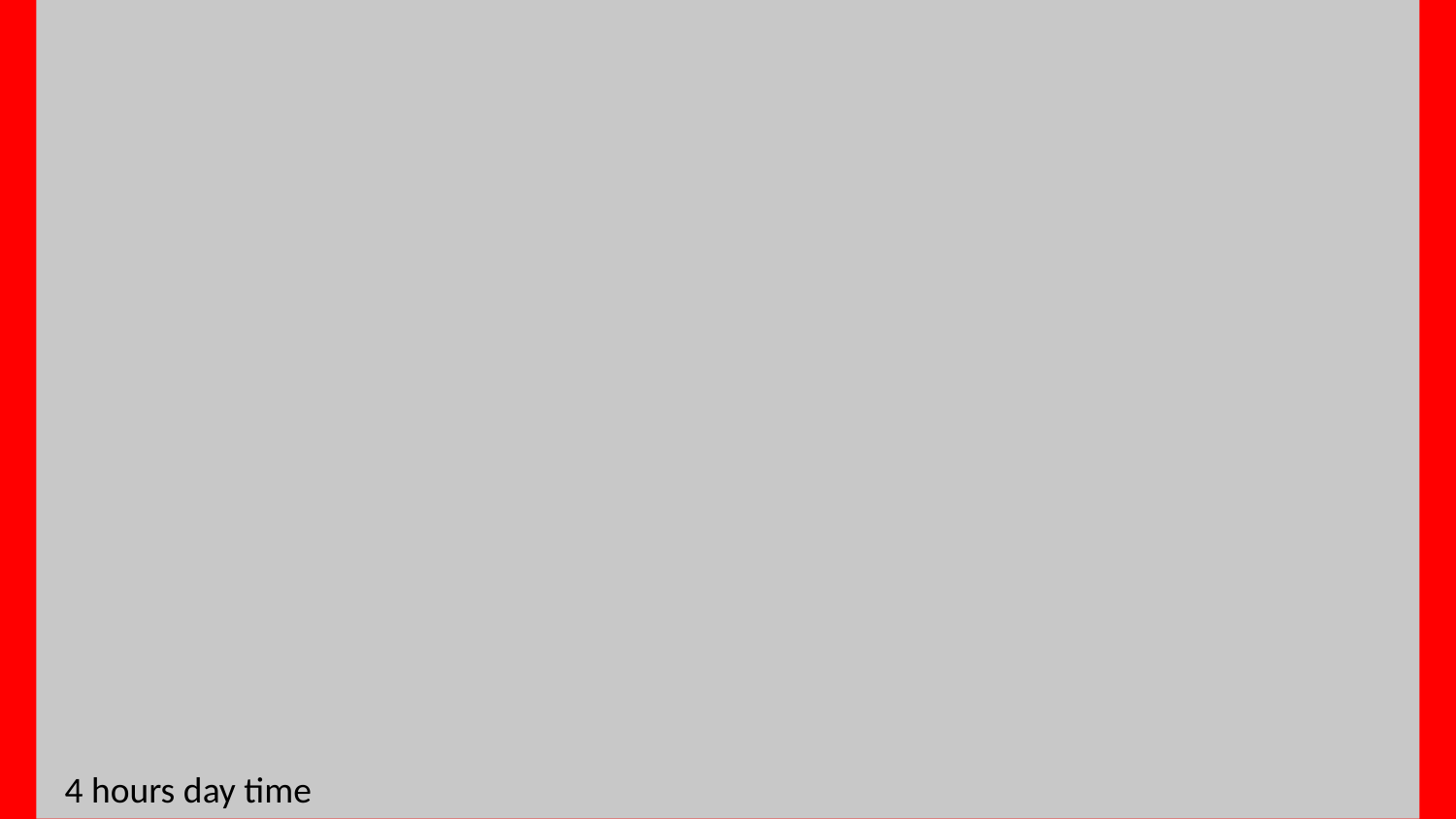

4 hours day time
