## Supplementary File 2 for "Combining supervised and unsupervised analyses to quantify behavioral phenotypes and validate therapeutic efficacy in a triple transgenic mouse model of Alzheimer’s disease": Supplementary_File_2.ijm

```
// Supplement S2. ImageJ and Fiji macro for automated analysis of 8 mouse cages
// This macro is designed for the automated analysis of mouse behavior

// To install the macro: open ImageJ, Plugins, install

// Macro 101022 original macro
// Macro 101122 adds aspect ratio and stretch-attend posture (SAP)
// Macro 101422 adds threshold 80, SAP ratio 2.3, only draw head if ratio > 2 (unreliable at low ratios)
// Macro 101722b adds frames per hour (fph) to correct hours and periods, removes unused ROI centroid, reorganized columns
// Macro 101722c uses fph for speed calculations, allows one to change the number of cages in the default settings
// Macro 101822 add dialog box, first crop then split channels to avoid memory issues on large files
// Macro 101922 major changes for large files: measure all cages in one duplicated image, then go to next in virtual stack
// Macro 101922b adds a calibration of the timer (needed for long recordings)
// Macro 102022 adds a step to skip images in the analysis
// Macro 102422 threshold 65, removes option to change cage number (errors)
// Macro 102822 Particle size >300 (without background subtraction)
// Macro 112322 threshold 40 to remove wet food from analysis
// Macro 120422 Adds dialog option for threshold and orientation of cage 1-8

// default settings

	ndraw = 8;				// number of measurements used for drawing centroid (green) and head (red)
	step = 1;				// can change this to reduce file size (would need to check speed measurements)
	ncages = 8;				// number of cages, keep at 8 to avoid ROI errors
	threshold = 40;			// 65 worked initially, but 40 is better when food gets wet at end of exp (exp 11/21/22)
							
// Works with ImageJ version 1.51j and up, earlier versions may work as well
	// requires("1.51j");

// Open video for image analysis
	waitForUser("Analysis of mouse behavior in 8-cage imaging system", 
	"Open video (File, Open) or images (File, Import, Image Sequence), Use virtual stack, OK, OK");	
	wait(100);
	dir=getDirectory("image");
	title=getTitle();
	rename("Video");
	Stack.getDimensions(wid, hei, cha, nimages, frames);

// Select fixed time point in video for calibration of timer (could add 12 hr mark in PowerPoint)
	waitForUser("Calibration of timer", 
	"If possible, move slider to the 12-hour point just before first visual stimuli (38,000-42,000 at 1 fps)");	
	wait(100);
	timemark = getSliceNumber();
	interval = 3600*12 / timemark;     // E.g. 3600 x 12 = 43,200 expected / 40,000 actual images = 1.08 sec

// DIALOG BOX to change the general default parameters

	Dialog.create("Analysis of mouse behavior");
	
	Dialog.addChoice("Cage1: ", newArray("window_down_right", "window_down_left", "window_up_right", "window_up_left"), "window_down_right");
	Dialog.addChoice("Cage2: ", newArray("window_down_right", "window_down_left", "window_up_right", "window_up_left"), "window_down_left");
	Dialog.addChoice("Cage3: ", newArray("window_down_right", "window_down_left", "window_up_right", "window_up_left"), "window_down_right");
	Dialog.addChoice("Cage4: ", newArray("window_down_right", "window_down_left", "window_up_right", "window_up_left"), "window_down_left");
	Dialog.addChoice("Cage5: ", newArray("window_down_right", "window_down_left", "window_up_right", "window_up_left"), "window_down_right");
	Dialog.addChoice("Cage6: ", newArray("window_down_right", "window_down_left", "window_up_right", "window_up_left"), "window_down_left");
	Dialog.addChoice("Cage7: ", newArray("window_down_right", "window_down_left", "window_up_right", "window_up_left"), "window_down_right");
	Dialog.addChoice("Cage8: ", newArray("window_down_right", "window_down_left", "window_up_right", "window_up_left"), "window_down_left");
	Dialog.addNumber("Number of images for analysis: ", nimages);
	Dialog.addNumber("* Interval between two images (sec): ", interval);
	Dialog.addNumber("Step (1 to analyze all images): ", step);	
	Dialog.addNumber("Threshold (40 for standard images, 50 for brighter images): ", threshold);	
	Dialog.addNumber("Number of measurements displayed on first image: ", ndraw);
	Dialog.addMessage("* Interval is calculated by the 12-hour calibration of the timer (change if needed)");
	Dialog.addMessage("  Interval ~ 1.06 at 1 fps with a 6% camera delay");
	
	Dialog.show();

	cage1window = Dialog.getChoice();
	cage2window = Dialog.getChoice();
	cage3window = Dialog.getChoice();
	cage4window = Dialog.getChoice();
	cage5window = Dialog.getChoice();
	cage6window = Dialog.getChoice();
	cage7window = Dialog.getChoice();
	cage8window = Dialog.getChoice();
	nimages = Dialog.getNumber();
	interval = Dialog.getNumber();	
	step = Dialog.getNumber();
	threshold = Dialog.getNumber();
	ndraw = Dialog.getNumber();
	
	wait(100);								// not sure: needed after dialog box?
	
	fph = 3600 / (interval * step);   			// actual frames per hour, e.g. 3600 / 1.08 = 3,333 fph
	
// Select ROIs
	setTool("rectangle");

	// ROI 1
	makeRectangle(71, 90, 240, 140);						// x, y, width, height
	waitForUser("Adjust ROI to select cage 1 (white bottom, not gray walls).  THEN click OK");		
	wait(100); 	
	getBoundingRect(Xroi1, Yroi1, Wroi1, Hroi1);			// stores ROI	

	// ROI 2
	makeRectangle(329, 88, 240, 140);
	waitForUser("Adjust ROI to select cage 2 (white bottom, not gray walls).  THEN click OK");		
	wait(100); 	
	getBoundingRect(Xroi2, Yroi2, Wroi2, Hroi2);			// stores ROI

	// ROI 3
	makeRectangle(73, 267, 240, 140);
	waitForUser("Adjust ROI to select cage 3 (white bottom, not gray walls).  THEN click OK");		
	wait(100); 	
	getBoundingRect(Xroi3, Yroi3, Wroi3, Hroi3);			// stores ROI

	// ROI 4
	makeRectangle(328, 264, 240, 140);
	waitForUser("Adjust ROI to select cage 4 (white bottom, not gray walls).  THEN click OK");		
	wait(100); 	
	getBoundingRect(Xroi4, Yroi4, Wroi4, Hroi4);			// stores ROI	

	// ROI 5
	makeRectangle(76, 559, 240, 140);
	waitForUser("Adjust ROI to select cage 5 (white bottom, not gray walls).  THEN click OK");		
	wait(100); 	
	getBoundingRect(Xroi5, Yroi5, Wroi5, Hroi5);			// stores ROI

	// ROI 6
	makeRectangle(331, 559, 240, 140);
	waitForUser("Adjust ROI to select cage 6 (white bottom, not gray walls).  THEN click OK");		
	wait(100); 	
	getBoundingRect(Xroi6, Yroi6, Wroi6, Hroi6);			// stores ROI

	// ROI 7
	makeRectangle(73, 731, 240, 140);
	waitForUser("Adjust ROI to select cage 7 (white bottom, not gray walls).  THEN click OK");		
	wait(100); 	
	getBoundingRect(Xroi7, Yroi7, Wroi7, Hroi7);			// stores ROI

	// ROI 8
	makeRectangle(331, 729, 240, 140);
	waitForUser("Adjust ROI to select cage 8 (white bottom, not gray walls).  THEN click OK");		
	wait(100); 	
	getBoundingRect(Xroi8, Yroi8, Wroi8, Hroi8);			// stores ROI

	getDateAndTime(year1, month1, dayOfWeek1, dayOfMonth1, hour1, minute1, second1, msec1);

// Save first image for drawing

	selectWindow("Video");	
	run("Select None");
	setSlice(1);	
	run("Duplicate...", "title=First_Image");    			// First image is later used to draw centroid and head position

// Duplicate one image, split channels, and MEASURE particles, then go to next image (allows for large files)

	setBatchMode(true);
	image = 1;

	run("Set Measurements...", "area mean centroid bounding fit display redirect=None decimal=1");

	while (image<=nimages) {                        		// loop for all images
		
	selectWindow("Video");
	setSlice(image);
	run("Duplicate...", "title=Image");			 			// sets aside a single image for analysis
	
	run("Split Channels");									// Only keep red channel
	close();
	close();
	selectWindow("Image (red)");
	rename(image);											// put the image number as a Label in the Results
	
	run("Gaussian Blur...", "sigma=3 stack");  				// blurs smaller objects and tail
	setThreshold(0, threshold, "raw");						// threshold is set in default settings (40)
	
	makeRectangle(Xroi1, Yroi1, Wroi1, Hroi1);
	run("Analyze Particles...", "size=300-Infinity display stack");	
	
	makeRectangle(Xroi2, Yroi2, Wroi2, Hroi2);
	run("Analyze Particles...", "size=300-Infinity display stack");	
	
	makeRectangle(Xroi3, Yroi3, Wroi3, Hroi3);
	run("Analyze Particles...", "size=300-Infinity display stack");
	
	makeRectangle(Xroi4, Yroi4, Wroi4, Hroi4);
	run("Analyze Particles...", "size=300-Infinity display stack");
	
	makeRectangle(Xroi5, Yroi5, Wroi5, Hroi5);
	run("Analyze Particles...", "size=300-Infinity display stack");
	
	makeRectangle(Xroi6, Yroi6, Wroi6, Hroi6);
	run("Analyze Particles...", "size=300-Infinity display stack");
	
	makeRectangle(Xroi7, Yroi7, Wroi7, Hroi7);
	run("Analyze Particles...", "size=300-Infinity display stack");
	
	makeRectangle(Xroi8, Yroi8, Wroi8, Hroi8);
	run("Analyze Particles...", "size=300-Infinity display stack");

	close();
	
	image = image + step;				// step1 analyzes every image, step6 analyzes image 1, 7, 13, 19 ...
	
	}                                    								// end of loop for all images

	setBatchMode(false);

// ADD CALCULATIONS IN RESULTS FILE

// Calculate position of the head 30 pixels away from centroid, along an angle

	row=0;

	while (row<nResults) {                        // loop for all rows in Results file
	Xcentroid = getResult ("X", row);	
	Ycentroid = getResult ("Y", row);
	Angle = getResult ("Angle", row);
	Radians = Angle * PI / 180;                  // sin and cos in ImageJ use radians
	Xhead1 = Xcentroid + 30*cos(Radians);        // two possible head positions along angle
	Yhead1 = Ycentroid - 30*sin(Radians);
	Xhead2 = Xcentroid - 30*cos(Radians);
	Yhead2 = Ycentroid + 30*sin(Radians);

	BX = getResult ("BX", row);                  // BX, BY is the left top of the Bounding Box
	BY = getResult ("BY", row);
	Width = getResult ("Width", row);
	Height = getResult ("Height", row);
	Xbox = BX + 0.5*Width;                       //  Xbox, Ybox is the center of the bounding box, towards head region
	Ybox = BY + 0.5*Height;

	BoxtoH1 = sqrt(pow(Xbox - Xhead1, 2) + pow(Ybox - Yhead1, 2));    // distance between center of box and possible head regions
	BoxtoH2 = sqrt(pow(Xbox - Xhead2, 2) + pow(Ybox - Yhead2, 2));

	if(BoxtoH1<BoxtoH2){
		Xhead = Xhead1;                         //  The region close to the center of box is set as the head region
		Yhead = Yhead1;
		}
	if(BoxtoH2<BoxtoH1){
		Xhead = Xhead2;
		Yhead = Yhead2;
		}	

	setResult("Xhead", row, Xhead);             // Adds values in the Results table
	setResult("Yhead", row, Yhead);

	row++;                              		// row +1 to go to next row
	}                                    		// end of loop for all rows

	updateResults();

// Add cage number

	row=0;
	cage = 0;

	while (row<nResults) {                        		// loop for all rows in Results file

	X = getResult ("X", row);	
	Y = getResult ("Y", row);

	if (X > Xroi1 && X < Xroi1+Wroi1 && Y > Yroi1 && Y < Yroi1+Hroi1) {cage = 1;}
	if (X > Xroi2 && X < Xroi2+Wroi2 && Y > Yroi2 && Y < Yroi2+Hroi2) {cage = 2;}	
	if (X > Xroi3 && X < Xroi3+Wroi3 && Y > Yroi3 && Y < Yroi3+Hroi3) {cage = 3;}
	if (X > Xroi4 && X < Xroi4+Wroi4 && Y > Yroi4 && Y < Yroi4+Hroi4) {cage = 4;}
	if (X > Xroi5 && X < Xroi5+Wroi5 && Y > Yroi5 && Y < Yroi5+Hroi5) {cage = 5;}
	if (X > Xroi6 && X < Xroi6+Wroi6 && Y > Yroi6 && Y < Yroi6+Hroi6) {cage = 6;}
	if (X > Xroi7 && X < Xroi7+Wroi7 && Y > Yroi7 && Y < Yroi7+Hroi7) {cage = 7;}
	if (X > Xroi8 && X < Xroi8+Wroi8 && Y > Yroi8 && Y < Yroi8+Hroi8) {cage = 8;}
	
	setResult("Cage", row, cage);            			// Adds values in the Results table
	row++;
	}														// end of rows
	
	updateResults();

// Add hours and periods of imaging. The frames per hour (fph) is set in the default settings

	row=0;

	while (row<nResults) {                        	// loop for all rows in Results file
		slice = getResultString("Label", row);	
		hour = Math.ceil (slice / fph);				// rounds up, eg slice 60 / 600 is 0.1 and rounds up to hour 1
		fpp = fph / 6;								// frames per period with 6 x 10-min periods per hour
		period = Math.ceil (slice / fpp);			// rounds up, eg slice 10 / 100 is 0.1 and rounds up to period 1 
		
		setResult("Hour", row, hour);
		setResult("Period", row, period);
		row++;                              		// row +1 to go to next row
		}                                    		// end of loop for all rows

	updateResults();

// Calculate quadrant of cage: A) left top, B) right top, C) left bottom, D) right bottom, using size of first ROI

	row=0;
	
	XCroi1 = Xroi1 + 0.5*Wroi1;							// Calculate center of ROI (X,Y)
	YCroi1 = Yroi1 + 0.5*Hroi1;
	XCroi2 = Xroi2 + 0.5*Wroi2;	
	YCroi2 = Yroi2 + 0.5*Hroi2;
	XCroi3 = Xroi3 + 0.5*Wroi3;	
	YCroi3 = Yroi3 + 0.5*Hroi3;
	XCroi4 = Xroi4 + 0.5*Wroi4;	
	YCroi4 = Yroi4 + 0.5*Hroi4;
	XCroi5 = Xroi5 + 0.5*Wroi5;	
	YCroi5 = Yroi5 + 0.5*Hroi5;
	XCroi6 = Xroi6 + 0.5*Wroi6;	
	YCroi6 = Yroi6 + 0.5*Hroi6;
	XCroi7 = Xroi7 + 0.5*Wroi7;	
	YCroi7 = Yroi7 + 0.5*Hroi7;
	XCroi8 = Xroi8 + 0.5*Wroi8;	
	YCroi8 = Yroi8 + 0.5*Hroi8;

	while (row<nResults) {     								// loop for all rows in Results file
	X = getResult ("X", row);	
	Y = getResult ("Y", row);
	cage = getResult ("Cage", row);

 	if(cage==1 && X<XCroi1 && Y<YCroi1){ quadrant = "A";}	// calculate if XY smaller or larger than the ROI center
 	if(cage==1 && X>XCroi1 && Y<YCroi1){ quadrant = "B";}	
 	if(cage==1 && X<XCroi1 && Y>YCroi1){ quadrant = "C";}	
 	if(cage==1 && X>XCroi1 && Y>YCroi1){ quadrant = "D";}	

 	if(cage==2 && X<XCroi2 && Y<YCroi2){ quadrant = "A";}	
 	if(cage==2 && X>XCroi2 && Y<YCroi2){ quadrant = "B";}	
 	if(cage==2 && X<XCroi2 && Y>YCroi2){ quadrant = "C";}	
 	if(cage==2 && X>XCroi2 && Y>YCroi2){ quadrant = "D";}	

 	if(cage==3 && X<XCroi3 && Y<YCroi3){ quadrant = "A";}	
 	if(cage==3 && X>XCroi3 && Y<YCroi3){ quadrant = "B";}	
 	if(cage==3 && X<XCroi3 && Y>YCroi3){ quadrant = "C";}	
 	if(cage==3 && X>XCroi3 && Y>YCroi3){ quadrant = "D";}	

 	if(cage==4 && X<XCroi4 && Y<YCroi4){ quadrant = "A";}	
 	if(cage==4 && X>XCroi4 && Y<YCroi4){ quadrant = "B";}	
 	if(cage==4 && X<XCroi4 && Y>YCroi4){ quadrant = "C";}	
 	if(cage==4 && X>XCroi4 && Y>YCroi4){ quadrant = "D";}	

 	if(cage==5 && X<XCroi5 && Y<YCroi5){ quadrant = "A";}	
 	if(cage==5 && X>XCroi5 && Y<YCroi5){ quadrant = "B";}	
 	if(cage==5 && X<XCroi5 && Y>YCroi5){ quadrant = "C";}	
 	if(cage==5 && X>XCroi5 && Y>YCroi5){ quadrant = "D";}	

 	if(cage==6 && X<XCroi6 && Y<YCroi6){ quadrant = "A";}	
 	if(cage==6 && X>XCroi6 && Y<YCroi6){ quadrant = "B";}	
 	if(cage==6 && X<XCroi6 && Y>YCroi6){ quadrant = "C";}	
 	if(cage==6 && X>XCroi6 && Y>YCroi6){ quadrant = "D";}	

 	if(cage==7 && X<XCroi7 && Y<YCroi7){ quadrant = "A";}	
 	if(cage==7 && X>XCroi7 && Y<YCroi7){ quadrant = "B";}	
 	if(cage==7 && X<XCroi7 && Y>YCroi7){ quadrant = "C";}	
 	if(cage==7 && X>XCroi7 && Y>YCroi7){ quadrant = "D";}	

 	if(cage==8 && X<XCroi8 && Y<YCroi8){ quadrant = "A";}	
 	if(cage==8 && X>XCroi8 && Y<YCroi8){ quadrant = "B";}	
 	if(cage==8 && X<XCroi8 && Y>YCroi8){ quadrant = "C";}	
 	if(cage==8 && X>XCroi8 && Y>YCroi8){ quadrant = "D";}	

 	setResult("Quad", row, quadrant);         				// Adds values in the Results table

	row++;             
	} 
	
	updateResults();

// Assign area name for each quadrant

	row = 0;

	while (row<nResults) {                 			// loop for all rows in Results file

	quadrant = getResultString("Quad", row);		// use String to get letters from Results table
	cage = getResult("Cage", row);
	
	home = 0;										// default value is 0 
	food = 0;
	wall = 0;
	window = 0;

	if(cage==1) {cagewindow=cage1window;}			// finds the location of the window in a specific cage
	if(cage==2) {cagewindow=cage2window;}
	if(cage==3) {cagewindow=cage3window;}
	if(cage==4) {cagewindow=cage4window;}
	if(cage==5) {cagewindow=cage5window;}
	if(cage==6) {cagewindow=cage6window;}
	if(cage==7) {cagewindow=cage7window;}
	if(cage==8) {cagewindow=cage8window;}

	if(cagewindow=="window_down_right" && quadrant=="A") { home=1;}
	if(cagewindow=="window_down_right" && quadrant=="B") { wall=1;}
	if(cagewindow=="window_down_right" && quadrant=="C") { food=1;}
	if(cagewindow=="window_down_right" && quadrant=="D") { window=1;}

	if(cagewindow=="window_down_left" && quadrant=="A") { wall=1;}
	if(cagewindow=="window_down_left" && quadrant=="B") { home=1;}
	if(cagewindow=="window_down_left" && quadrant=="C") { window=1;}
	if(cagewindow=="window_down_left" && quadrant=="D") { food=1;}

	if(cagewindow=="window_up_right" && quadrant=="A") { food=1;}
	if(cagewindow=="window_up_right" && quadrant=="B") { window=1;}
	if(cagewindow=="window_up_right" && quadrant=="C") { home=1;}
	if(cagewindow=="window_up_right" && quadrant=="D") { wall=1;}

	if(cagewindow=="window_up_left" && quadrant=="A") { window=1;}
	if(cagewindow=="window_up_left" && quadrant=="B") { food=1;}
	if(cagewindow=="window_up_left" && quadrant=="C") { wall=1;}
	if(cagewindow=="window_up_left" && quadrant=="D") { home=1;}
	
	setResult("Home", row, home);
	setResult("Food", row, food);
	setResult("Wall", row, wall);
	setResult("Window", row, window);
	
	row++;                              		// row +1 to go to next row
	}                                    		// end of loop for all rows

	updateResults();

// In or out? Out: Wall or window area.

	row = 0;

	while (row<nResults) { 

		home = getResult("Home", row);
		food = getResult("Food", row);
		wall = getResult("Wall", row);
		window = getResult("Window", row);
		
		if(home == 1) {out=0;}
		if(food == 1) {out=0;}				
		if(wall == 1) {out=1;}
		if(window == 1) {out=1;}

	setResult("Out", row, out);
	row++;   
	}

	updateResults();

// Speed in cm per second

	row = 10;
	scale = 76/70;									// Scale: 76 mm mouse arc is 70 pix

	while (row<nResults) {                 			// loop for all rows in Results file

	imagecurr = getResult("Label", row);
	cagecurr = getResult("Cage", row);
	
	cageprev10 = getResult("Cage", row-10);			// looks for cage numbers in previous 10 rows
	cageprev9 = getResult("Cage", row-9);
	cageprev8 = getResult("Cage", row-8);
	cageprev7 = getResult("Cage", row-7);
	cageprev6 = getResult("Cage", row-6);
	cageprev5 = getResult("Cage", row-5);
	cageprev4 = getResult("Cage", row-4);
	cageprev3 = getResult("Cage", row-3);
	cageprev2 = getResult("Cage", row-2);
	cageprev1 = getResult("Cage", row-1);

	if (cagecurr == cageprev10){lastrow = 10;}		// finds last row with same cage number (e.g. 10 rows up)
	if (cagecurr == cageprev9){lastrow = 9;}
	if (cagecurr == cageprev8){lastrow = 8;}
	if (cagecurr == cageprev7){lastrow = 7;}
	if (cagecurr == cageprev6){lastrow = 6;}
	if (cagecurr == cageprev5){lastrow = 5;}
	if (cagecurr == cageprev4){lastrow = 4;}
	if (cagecurr == cageprev3){lastrow = 3;}
	if (cagecurr == cageprev2){lastrow = 2;}
	if (cagecurr == cageprev1){lastrow = 1;}

	Xcurr = getResult("X", row);	
	Ycurr = getResult("Y", row);	
	Xprev = getResult("X", row-lastrow);			// Gets XY of previous measurement in same cage
	Yprev = getResult("Y", row-lastrow);

	pixmoved = sqrt(pow(Xcurr - Xprev, 2) + pow(Ycurr - Yprev, 2));
	
	speedmmps = (pixmoved * scale) / (interval * step);	// speed in mm per sec
	speed = speedmmps / 10;								// speed in cm per sec

	if (imagecurr<=10) {speed = 0;}						// cannot calculate speed on first images
	
	if (speed<1) {move = 0;}	
	if (speed>1) {move = 1;}							// speed is more than 1 cm per second
	
	setResult("Speed", row, speed);
	setResult("Move", row, move);
	row++;  
	}

	updateResults();

// calculate aspect ratio (major / minor axis of elipse) and SAP (elongated without movement)

	row = 0;

	while (row<nResults) { 

		major = getResult("Major", row);
		minor = getResult("Minor", row);
		speed = getResult("Speed", row);		
		aspectratio = major / minor;

		sap = 0;											// stretch-attend posture (SAP)	
		if (speed<1 && aspectratio>2.3) {sap = 1;}			// eccentricity 0.9 = ratio of 2.3	

		setResult("Ratio", row, aspectratio);
		setResult("SAP", row, sap);
		row++;   
		}

	updateResults();
	
// DRAW CENTROID AND HEAD ON FIRST IMAGE

	row=0;
	selectWindow("First_Image");
	setLineWidth(10);
	
	if (ndraw>nResults){ndraw=nResults;}						// cannot draw more points than nResults

	while (row<ndraw) {        
		X=getResult ("X", row);
		Y=getResult ("Y", row);
		Xhead=getResult ("Xhead", row);	
		Yhead=getResult ("Yhead", row);	
		ratio = getResult ("Ratio", row);
		setColor(0, 255, 0);
		drawLine(X, Y, X, Y);                   				// drawLine works better than drawOval
		setColor(255, 0, 0);
		if(ratio>2) {drawLine(Xhead, Yhead, Xhead, Yhead);}  	// head unreliable when mouse is curled up

		row++;
		}              

// ADD INFO AT THE END OF THE RESULTS FILE

	setResult("Label", nResults, "    ");       		// can add + "\t" + "word" for a tab in the table
	setResult("Label", nResults, title);				// print title and directory
	setResult("Label", nResults, dir);					// nResults adds a row to the Results table

	setResult("Label", nResults, "   ");
	getDateAndTime(year2, month2, dayOfWeek2, dayOfMonth2, hour2, minute2, second2, msec2);
	setResult("Label", nResults, "Start of automated analysis:  " + month1+1 + "/" + dayOfMonth1 + "/" + year1 + "   time: " + hour1 + ":" + minute1 + ":" + second1);  // +1 needed in month?!
	setResult("Label", nResults, "End of automated analysis:  " + month2+1 + "/" + dayOfMonth2 + "/" + year2 + "   time: " + hour2 + ":" + minute2 + ":" + second2);

	setResult("Label", nResults, "   ");
	setResult("Label", nResults, "Threshold: " + threshold);
	setResult("Label", nResults, "Interval between two images (sec): " + interval);
	setResult("Label", nResults, "   ");
	
	setResult("Label", nResults, "R0I 1:  X=" + Xroi1 + ", Y=" + Yroi1 + ", Width="+ Wroi1 + ", Height="+ Hroi1);
	setResult("Label", nResults, "R0I 1:  X=" + Xroi1 + ", Y=" + Yroi1 + ", Width="+ Wroi1 + ", Height="+ Hroi1);
	setResult("Label", nResults, "R0I 3:  X=" + Xroi3 + ", Y=" + Yroi3 + ", Width="+ Wroi3 + ", Height="+ Hroi3);
	setResult("Label", nResults, "R0I 4:  X=" + Xroi4 + ", Y=" + Yroi4 + ", Width="+ Wroi4 + ", Height="+ Hroi4);
	setResult("Label", nResults, "R0I 5:  X=" + Xroi5 + ", Y=" + Yroi5 + ", Width="+ Wroi5 + ", Height="+ Hroi5);
	setResult("Label", nResults, "R0I 6:  X=" + Xroi6 + ", Y=" + Yroi6 + ", Width="+ Wroi6 + ", Height="+ Hroi6);
	setResult("Label", nResults, "R0I 7:  X=" + Xroi7 + ", Y=" + Yroi7 + ", Width="+ Wroi7 + ", Height="+ Hroi7);
	setResult("Label", nResults, "R0I 8:  X=" + Xroi8 + ", Y=" + Yroi8 + ", Width="+ Wroi8 + ", Height="+ Hroi8);
	setResult("Label", nResults, "   ");

	setResult("Label", nResults, "Cage1: " + cage1window);
	setResult("Label", nResults, "Cage2: " + cage2window);
	setResult("Label", nResults, "Cage3: " + cage3window);
	setResult("Label", nResults, "Cage4: " + cage4window);
	setResult("Label", nResults, "Cage5: " + cage5window);
	setResult("Label", nResults, "Cage6: " + cage6window);
	setResult("Label", nResults, "Cage7: " + cage7window);
	setResult("Label", nResults, "Cage8: " + cage8window);


// Automatically save results to Image directory with specified file name

	filename = "Results_" + title + ".xls";
	saveAs("Results", dir+filename);

	selectWindow("First_Image");
	imagename = "First_Image_" + title + ".jpg";
	saveAs("Jpeg", dir+imagename);

	beep();

// end of macro
```
